# Generative replay across hippocampal-neocortical circuits

**DOI:** 10.64898/2026.07.24.740539

**Authors:** Cal M. Shearer, Rosalind McDonald-Hill, Mathilde C.C. Guillaumin, Olivia J. McGinnis, Vitor Lopes-Dos-Santos, Pavel V. Perestenko, David Dupret, Helen C. Barron

## Abstract

Flexible behaviour requires predictions that extend beyond direct experience, yet the neural mechanisms that support this process remain unclear. A candidate mechanism occurs during hippocampal sharp-wave ripples (SWRs), in which generative replay combines representations for discrete past events into novel activity patterns. However, it remains unknown whether comparable computations occur in neocortex. Using simultaneous electrophysiology and calcium imaging across hippocampus and neocortex, we record neural activity in freely moving mice during an inference task and in subsequent sleep. We show that neuronal sequences in primary visual cortex (V1) encode relationships that were never directly experienced, providing evidence for generative replay outside the hippocampus. Generative replay in V1 is predicted by preceding hippocampal activity, suggesting that hippocampal activity provides a teaching signal to reorganise neocortical representations during sleep. These findings identify a hippocampal-neocortical mechanism for generative replay which may support the construction of a hierarchical generative model to enable flexible behaviour.

## Introduction

Memory consolidation can be defined as a process that stabilises and redistributes memories for long-term storage^1,2^. “Offline” periods of rest and sleep are thought to facilitate memory consolidation through neural computations occurring in the hippocampus during high frequency oscillations, termed Sharp-Wave Ripples (SWRs) or ripples^3–12^. During ripples, hippocampal spiking activity recapitulates or “replays” past experience on a temporally compressed scale^5,7–9,13,14^, a process thought to strengthen and stabilise memories by enabling synaptic plasticity. Consistent with this view, causal disruption of hippocampal ripples can impair subsequent memory performance^6,11,12^, while lengthening ripples can enhance performance^15^.

The hippocampus is anatomically well positioned to coordinate systems-level memory consolidation. Sitting at the apex of the cortical hierarchy, the hippocampus receives convergent inputs from and sends diverging projections back to distributed neural circuits. Replay events generated within the hippocampus may therefore propagate beyond the hippocampus. Indeed, empirical evidence suggests that hippocampal activity during ripples can predict activity in primary sensory cortices, including primary visual cortex^16^, primary auditory cortex^17^ and somatosensory cortex^18^, while coordinated activity is also observed in higher-order brain regions such as entorhinal cortex^19^ and prefrontal cortex^20^. These findings support the “two-stage” model of consolidation^1,2,8,21^, in which the hippocampus rapidly encodes new experiences before replay drives reactivation across neocortical circuits, ultimately stabilising and redistributing memories for long-term storage.

However, although replay has traditionally been viewed as a faithful recapitulation of past experience, hippocampal replay can extend beyond direct experience^22–28^ to provide an example of “generative replay”. This generative replay is thought to reflect active sampling of an internal model of the world^29–31^ and is thought to provide powerful computational solutions to multiple aspects of higher-order cognition^29–33^. For example, by co-reactivating memories acquired at different time points^22,28^, generative replay can establish new associations and relationships to facilitate novel predictions and planning^26,27^, inference^22^ and generalisation^34^, while also enabling credit assignment^28,35^. Thus, beyond consolidating existing memories, generative replay provides a candidate mechanism to *reorganise* the content of memory to support flexible cognition.

Critically, it remains unknown whether generative replay in the hippocampus influences activity in neocortical circuits. If hippocampal generative replay provides a training signal to neocortex, this may endow neocortical circuits with capacity to build and consolidate neural representations that deviate from past-experience. Rather than simply strengthening existing memories, hippocampal replay may therefore reorganise the intrinsic *content* of memories, not only in the hippocampus but across the entire cortical hierarchy. This would implicate hippocampal generative replay as a mechanism with capacity to build a deep *hierarchical generative model*^30,31,36^ that supports novel predictions via abstraction, inference, and generalization—the hallmarks of flexible and adaptive behaviour.

This framework gives rise to three predictions. First, generative sequences initiated in the hippocampus should propagate to sensory cortex. Second, generative activity in sensory cortex should occur after that observed in the hippocampus. Third, with learning, generative replay in sensory cortex should become progressively less dependent on hippocampal input, reflecting the consolidation of generative representations in neocortex.

Here we sought to test these predictions by investigating generative replay across hippocampal-neocortical networks. To this end, we implement a multi-day inference task in mice^22^. The task includes an *inference test* where animals are required to make inferential decisions that extend beyond direct experience. Thus, the task is designed to encourage animals to reorganise mnemonic content to create new knowledge that facilitates future decisions. By combining multi-unit electrophysiology and calcium imaging across hippocampal-neocortical circuits, we investigate the interaction between hippocampus and neocortical circuits during the inference task and in subsequent periods of sleep. Specifically, we ask whether hippocampal-neocortical circuits engage in generative replay during sleep, to build a hierarchical generative model.

## Results

### Mouse behavioural performance on the inference task

To investigate generative replay across hippocampal-neocortical circuits, we leveraged an inference task previously used to record brain activity during inferential choice in both humans and mice^22^. The inference task included 3 stages and was performed in mice over the course of 18 days, in an open-field environment (Figure 1a-e). In the first stage (“Observational learning”, Figure 1a), mice learned to associate an auditory cue (*X_n)_* with a corresponding visual cue (*Y_n)_*, for two sets of cues (set 1 and set 2). In the second stage (“Conditioning”, Figure 1a), mice learned to associate the previously seen visual cues with either a rewarding outcome (set 1, sucrose, *Z_1)_* or a neutral outcome (set 2, water, *Z_2)_*, delivered to a dispenser in the open field (Figure 1b). In the final stage (“Inference test”, Figure 1a), we tested whether the mice could infer an association between the auditory cue (*X_n)_* and the outcome (*Z_n)_* despite these two cues never being directly experienced together. Thus, the mice experienced *X_n_* → *Y_n_* and *Y_n_* → *Z_n,_* but never *X_n_* → *Z_n._* Importantly, this inference task provides opportunity to investigate how new inferred relationships are constructed across hippocampal-neocortical circuits.

**Figure 1.**
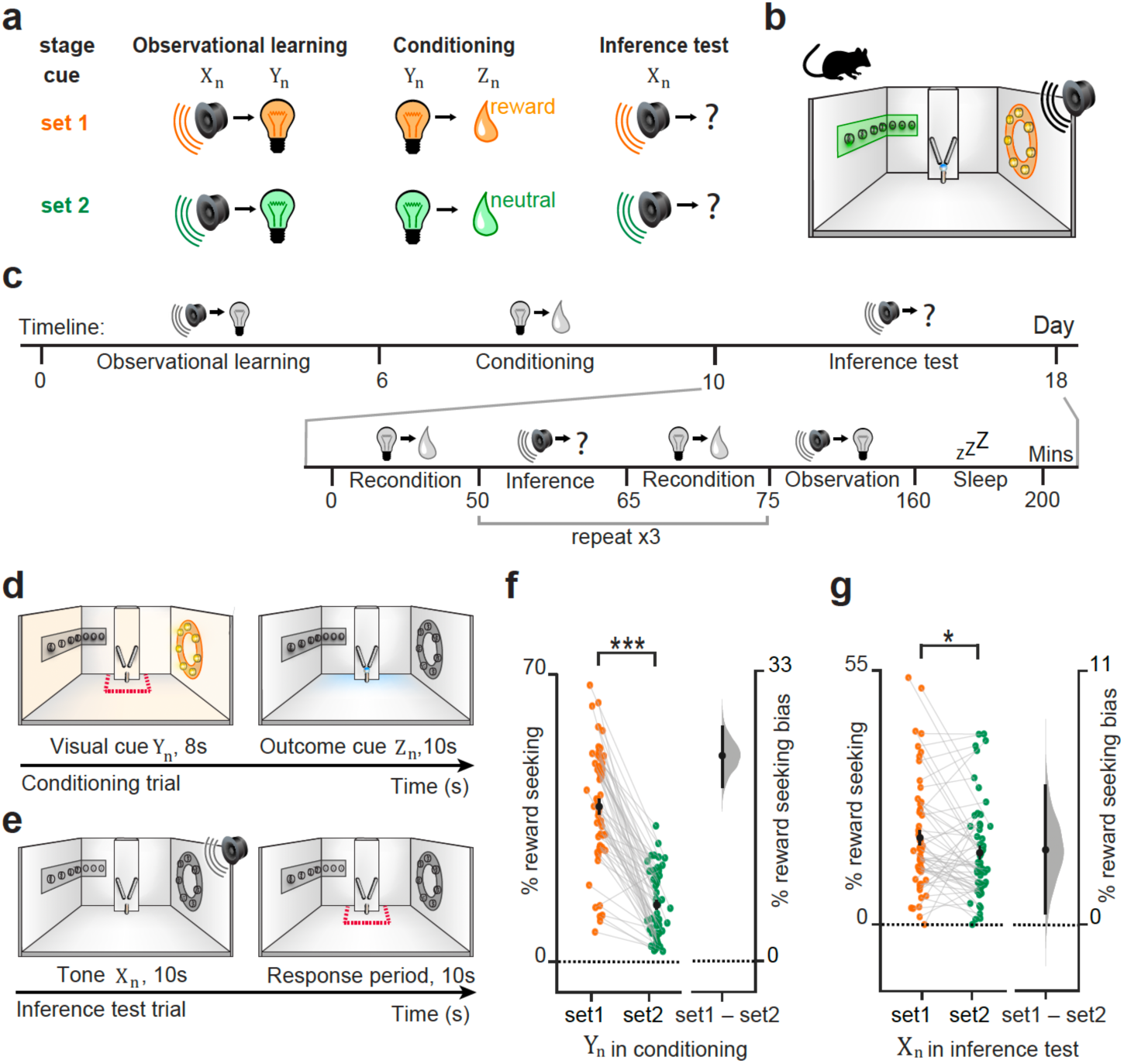
Inference task protocol and behavioural performance in mice. **(a)** Schematic showing the overall structure of the three-stage inference task. Mice first learned to associate an auditory cue *(X_n)_* with a visual cue *(Y_n)_* (*Observational Learning*). Next, they learned to associate a visual cue *(Y_n)_* with an outcome *(Z_n)_ (Conditioning)*, where the outcome was rewarding in set 1 (orange) and neutral in set 2 (green). In the *Inference test*, an auditory cue *(X_n)_* was presented in isolation and reward-seeking behaviour quantified as a measure of inference from *X_n_* to *Z_n._* **(b)** Schematic showing the open field arena in which the mice performed the task. Auditory cues were played into the environment using a speaker positioned above the open field. Visual cues were presented using two different LED strips on the walls of the environment. Outcome cues were delivered to a liquid dispenser positioned against one wall. **(c)** Timeline of the three-stage inference task, in days, with details of an example recording day, in minutes. **(d-e)** Schematic showing an example trial in the *Conditioning* stage (*d*) and in the *Inference test* (*e*), performed within the open field. Reward seeking behaviour was quantified using time spent in the area around the dispenser, indicated by the red dotted line, during the visual cue *(Y_n)_* (*Conditioning* trials), or after the auditory cue *(X_n)_* when no cue is presented (*Inference test* trials). **(f-g)** Left of each panel: raw data points for set 1 (orange) and set 2 (green). Each data point represents one recording day and shows the average proportion of time spent in the area around the dispenser (indicated by red dotted line, *d-e*), during the visual cue (*Y_n)_* in *Conditioning* trials (*f*), or immediately following the auditory cue (*X_n)_* in the *Inference test* trials (*g*). Right of each panel: difference in means between set 1 (orange) and set 2 (green) shown using bootstrap-coupled estimation (DABEST) plots^37^. Effect size for the difference between set 1 and 2, computed from 10,000 bias-corrected bootstrapped resamples^38^: black dot, mean; black ticks, 95% confidence interval; filled-curve, sampling-error distribution. Mice showed significant reward-seeking bias to cues in set 1 (higher proportion of time spent in the area around the dispenser in set 1 relative to set 2) during visual cues (*Y_n)_* in *Conditioning* (p < 0.001; *f*) and immediately following auditory cues (*X_n)_* in the *Inference test* (p=0.015; *g*). Panels *a-e* adapted from^22^.

To first assess behavioural evidence for learning and inference, we quantified reward-seeking behaviour, during both the conditioning (*Y_n_* to *Z_n)_* and inference (*X_n_* in isolation) stages of the task respectively. Reward-seeking behaviour was defined as the amount of time spent in the area around the dispenser, either during the LED (*Y_n_* in conditioning trials, Figure 1d) or in the 10 s period after the tone (*X_n_* in the inference test, Figure 1e). We expected to see higher levels of reward-seeking behaviour in response to cues in set 1, the set that included the rewarding outcome (*Z_1)_*, compared to cues in set 2, the set that included the neutral outcome (*Z_2)_*.

To confirm that mice were successfully conditioned, we quantified reward seeking behaviour on “reconditioning” trials (Figure 1c), which were presented immediately prior to each inference test. As expected, during reconditioning trials, mice showed higher levels of reward-seeking behaviour during visual cues in set 1 (*Y_1)_* relative to set 2 (*Y_2)_* (*Y_1_* vs. *Y_2,_* Figure 1f). Therefore, mice spent a greater proportion of time by the dispenser during the LED associated with sucrose (*Y_1)_* compared to the LED associated with water (*Y_2)_*, prior to outcome delivery. This demonstrates that the mice successfully learned to associate the visual cues (*Y_n)_* with the respective outcomes (*Z_n)_*.

During the inference test, we asked whether the mice could successfully infer an association between the auditory cues (*X_n)_* and the outcomes (*Z_n)_*, even though these cues had not been directly experienced together and the outcomes (*Z_n)_* were never presented during the inference test. In response to the auditory cues (*X_n)_*, mice showed significantly higher levels of reward-seeking behaviour in response to cues in set 1 (*Z_1)_* relative to set 2 (*Z_2)_* (Figure 1g). This demonstrates that mice were able to infer a relationship between the auditory cues and outcomes (*X_n_* → *Z_n)_*, even though they never experienced these cues together, replicating previous results using this inference task^22^.

### Anatomical pathway between dCA1 and V1

Next, we aimed to establish hippocampal-neocortical interactions during both task performance and in subsequent periods of sleep. We hypothesised that interactions between hippocampus and neocortex during sleep may allow generative replay in the hippocampus to reach neocortical circuits, to build mnemonic representations across the cortical hierarchy that extend beyond direct experience.

Previous neuroimaging studies using the same inference task in humans suggest that correct inferential choice is supported by a network of brain regions involved in both memory and the processing of relevant sensory cues, namely hippocampus, retrosplenial cortex (RSC) and primary visual cortex (V1)^22^. This raises the possibility that computations that support performance on the inference task are mediated by activity across a hippocampus-RSC-V1 anatomical pathway, where hippocampal generative replay can influence computations in RSC and V1.

To test this hypothesis, we first characterised the anatomical pathway between hippocampus, RSC and V1. While there are no direct anatomical connections between hippocampus and V1, previous studies in mice have identified RSC as a candidate intermediary area^39–42^. However, it remains unclear whether there is a disynaptic pathway from hippocampus, to RSC, to V1, such that neurons in RSC receiving dCA1 input also project to V1 (Figure 2a).

**Figure 2.**
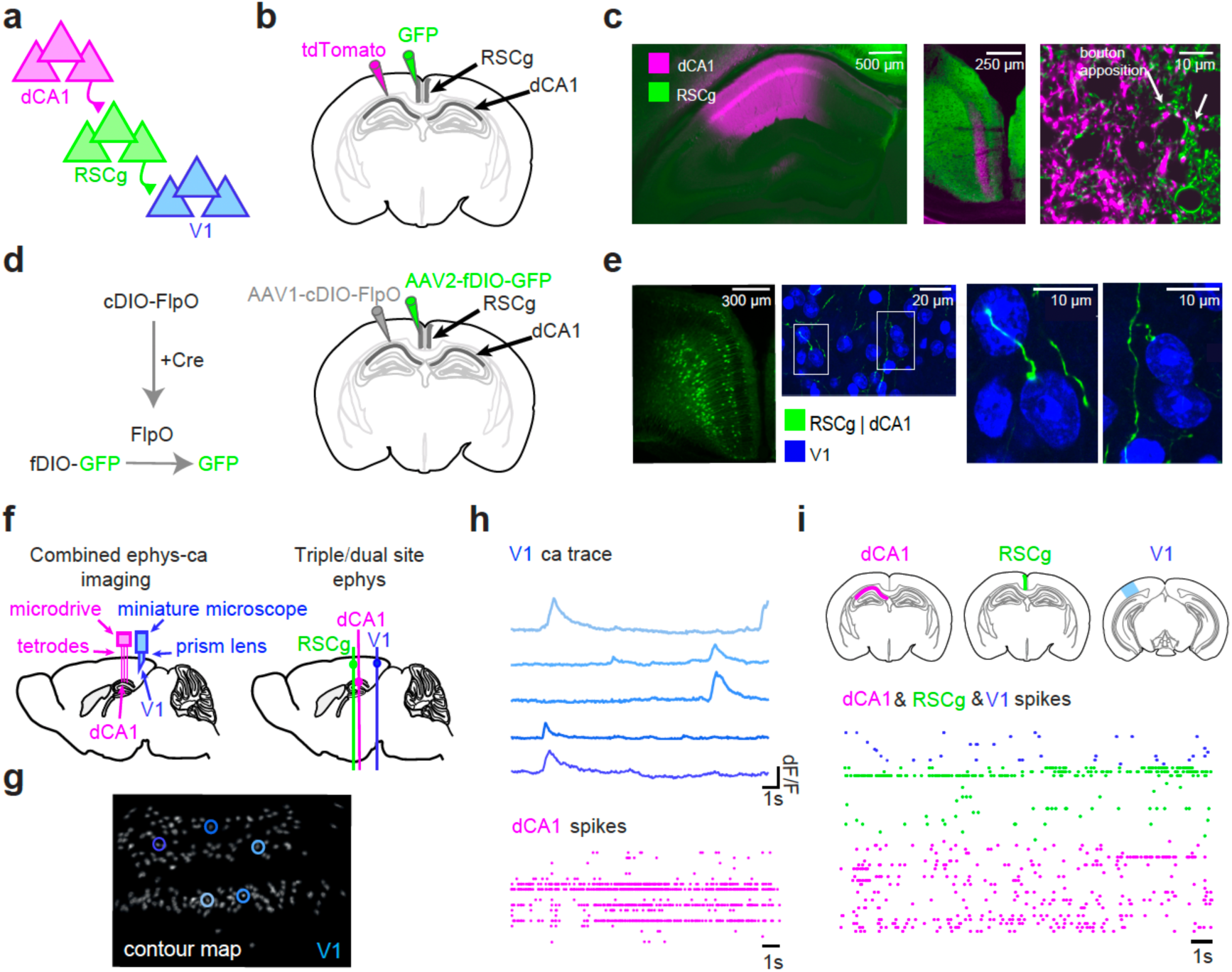
Anatomical pathway from dCA1 to V1 and experimental set up for recording in-vivo neuronal activity. **(a)** Schematic showing proposed anatomical pathway from dCA1 to V1 via RSCg. **(b)** Schematic showing injection sites in a CaMKIIα-Cre mouse, used to transduce dCA1 principal cells with Cre-dependent tdTomato (pink) viral construct and RSCg neurons with GFP viral construct (green). **(c)** tdTomato and GFP expression in dCA1 and RSC, respectively, showing dCA1 axonal projections (in pink) to RSCg (green). Hippocampus (left); RSC (middle); RSCg neurons (green) with putative bouton apposition (pink) from dCA1 axonal projections (right). **(d)** Schematic showing trans-synaptic intersectional strategy in a CAMKIIα-Cre mouse to restrict the expression of GFP to excitatory RSCg cells receiving dCA1 input. Left: Translation of fDIO-GFP construct is conditional to the sequential activity of Cre and FlpO recombinases, adapted from Trouche et al.^43^. Right: This strategy was used for trans-synaptic anterograde targeting of dCA1-connected RSCg cells. AAV1-cDIO-FlpO was injected into dCA1 and AAV2-fDIO-GFP was injected into RSCg. GFP is only expressed in the presence of both viruses, in excitatory cells expressing cre. **(e)** Left: RSC section with cells labelled using the trans-synaptic viral strategy. Middle: Example V1 soma (in blue, following DAPI staining for cell nuclei) with putative axonal targeting from RSCg neurons that receive dCA1 input (green). Two white boxes indicate regions of interest, shown on *Right*. Right: Two examples of V1 soma (in blue, following DAPI staining for cell nuclei) with putative contacts from RSCg axonal projections (green), where RSCg labelled neurons receive dCA1 input. **(f)** Left: Schematic showing simultaneous recording of electrophysiology data in dCA1 and calcium imaging data from V1 (n=4). Right: Schematic showing simultaneous recording of electrophysiology data across multiple brain regions, to allow for triple (n=2) or dual (n=3) site recording across dCA1, RSCg and/or V1. **(g)** Example contour map for cells imaged in V1 using combined electrophysiology and calcium imaging set up shown in *f*. **(h)** Example data recorded using the combined electrophysiology and calcium imaging set-up shown in *f*, with calcium traces from V1 (top, blue) and spikes from dCA1 (bottom, pink). **(i)** Example data recorded using triple site electrophysiology set up shown in *f*, with spikes recorded from V1 (top, blue), RSCg (middle, green) and dCA1 (bottom, pink).

**Supplementary Figure 1.**
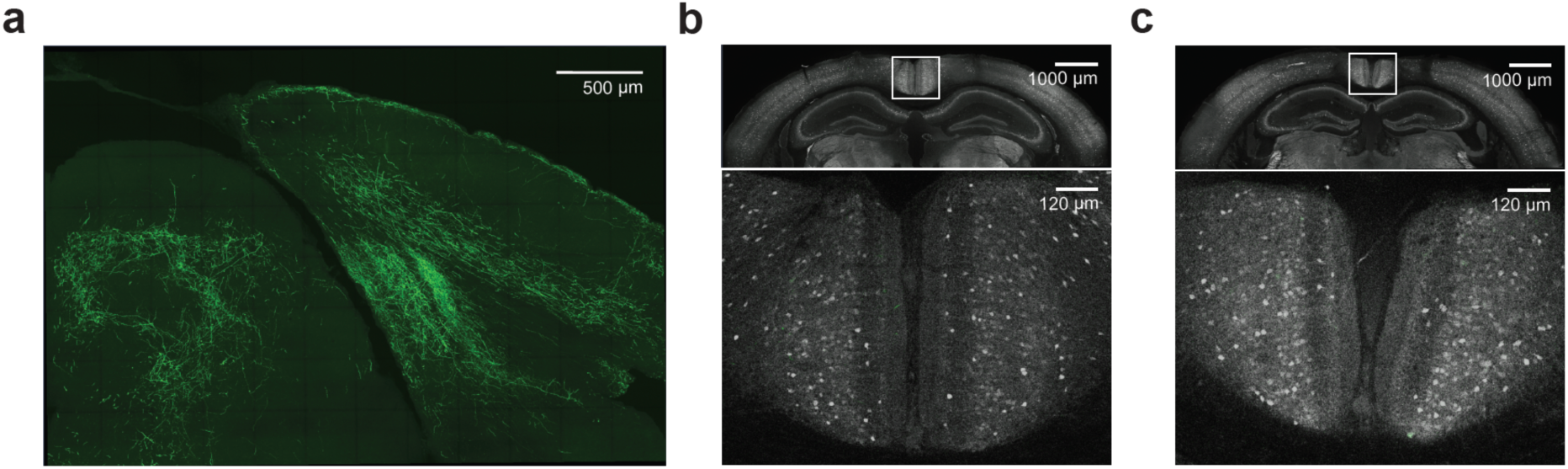
Anatomical pathway from dCA1 to V1, supporting. Figure 2 **(a)** Confocal image from the same mouse as shown in Figure 2e, showing V1 with axonal projections in green that derive from excitatory RSCg cells that receive dCA1 input (imaged in GFP-detection range only). **(b-c)** Controls for trans-synaptic virus used in Figure 2d-e. **(b)** Control for non-specific expression due to pipette-leakage: In this control, the first construct (EF1a-DIO-FLPo-Myc) was lowered into dCA1 via pipette but retracted without injection to assess potential leakage from the pipette track. The second construct (EF1a-FDIO-EGFP) was injected into the RSCg as normal. *Upper:* Confocal image showing full span of CamKII-cre mouse brain, in both the GFP and Dylight 405 detecting channels. The latter channel labels parvalbumin positive (PV+) interneurons and is shown for reference. *Lower:* Zoomed in view of the RSCg, showing an absence of GFP signal despite imaging in the GFP detection range. **(c)** Control for spontaneous recombination: In this control, the first construct (targeting dCA1) was omitted, and only the second construct (EF1a-FDIO-EGFP) was injected into the RSCg to confirm the necessity of the first virus for GFP expression. *Upper:* Confocal image showing full span of control CamKII-cre mouse brain in both the GFP and Dylight 405 detecting channels. The latter channel labels PV+ interneurons and is shown for reference. *Lower:* Zoomed in view of the RSCg, showing negligible GFP signal despite imaging in the GFP detection range.

To investigate the candidate dCA1-RSCg-V1 pathway, we first traced the pathway between dCA1 and RSC. We transduced dCA1 pyramidal neurons in CamKII-Cre mice with a Cre-dependent tdTomato viral construct, and neurons in granular RSC (RSCg) with a Green Fluorescent Protein (GFP) viral construct (Figure 2b). We then acquired confocal images of RSCg sections and saw evidence for putative bouton apposition between axons from pyramidal cells in dCA1 and neuronal cell bodies in RSCg (Figure 2c), suggesting that dCA1 pyramidal neurons project to neurons in RSCg.

Next, we asked whether some of the dCA1 neurons that project to RSCg provide the first synapse in a disynaptic pathway from dCA1 to V1. To test this we used a trans-synaptic intersectional strategy which allowed us to selectively express GFP in RSCg pyramidal neurons receiving monosynaptic inputs from dCA1 (Figure 2d)^43^. This approach was based on a double-conditional Boolean logic of transgene expression, where GFP is expressed in presence of both Cre and flippase (FlpO) recombinases which act in series within the same cell. After implementing this trans-synaptic intersectional strategy, we acquired confocal images of V1 sections that had been counterstained with DAPI. In V1, we saw evidence for putative bouton apposition between axons deriving from RSCg (labelled with GFP) and neuronal cell bodies in V1 (Figure 2e, Supplementary Figure 1). This suggests evidence for a disynaptic pathway from dCA1 to V1 via RSCg, where neurons in RSCg that receive inputs from dCA1 project to V1 (Figure 2a).

Having established a putative anatomical pathway from dCA1 to V1 via RSCg, we targeted these regions for in vivo recordings of neuronal activity during the inference task using two approaches (Figure 2f). First, we combined calcium imaging of V1 cells with electrophysiological recordings in dCA1 (Figure 2f-h). This combined approach provided a high cell yield in V1, while allowing recording of both the local field potential (LFP) and spiking activity in dCA1. To also record from RSCg, we employed triple and dual site electrophysiological recordings targeting dCA1, RSC, and/or V1 (Figure 2f, 2i), providing high temporal resolution recordings across all three brain regions.

### V1 represents a prospective code during inference

Having characterised a disynaptic anatomical pathway from dCA1→RSC→V1, we hypothesised that activity in both dCA1 and V1 should support inferential choice, as observed in humans^22^. To test this, we trained a cross-validated logistic regression classifier to predict inferential choice. Thus, we used neural activity recorded during the auditory cues (*X_n)_* to predict behavioural data (Figure 1g), where we quantified the accuracy of inferential choice according to the presented cue using visits to the reward dispenser after the tone in the inference test. We were able to significantly predict inferential choice from neural activity in both dCA1 (mean accuracy 71.64%) and V1 (mean accuracy 66.98%) (Figure 3a). This suggests that information about inferential choice is represented by neurons in both dCA1 and V1.

**Figure 3.**
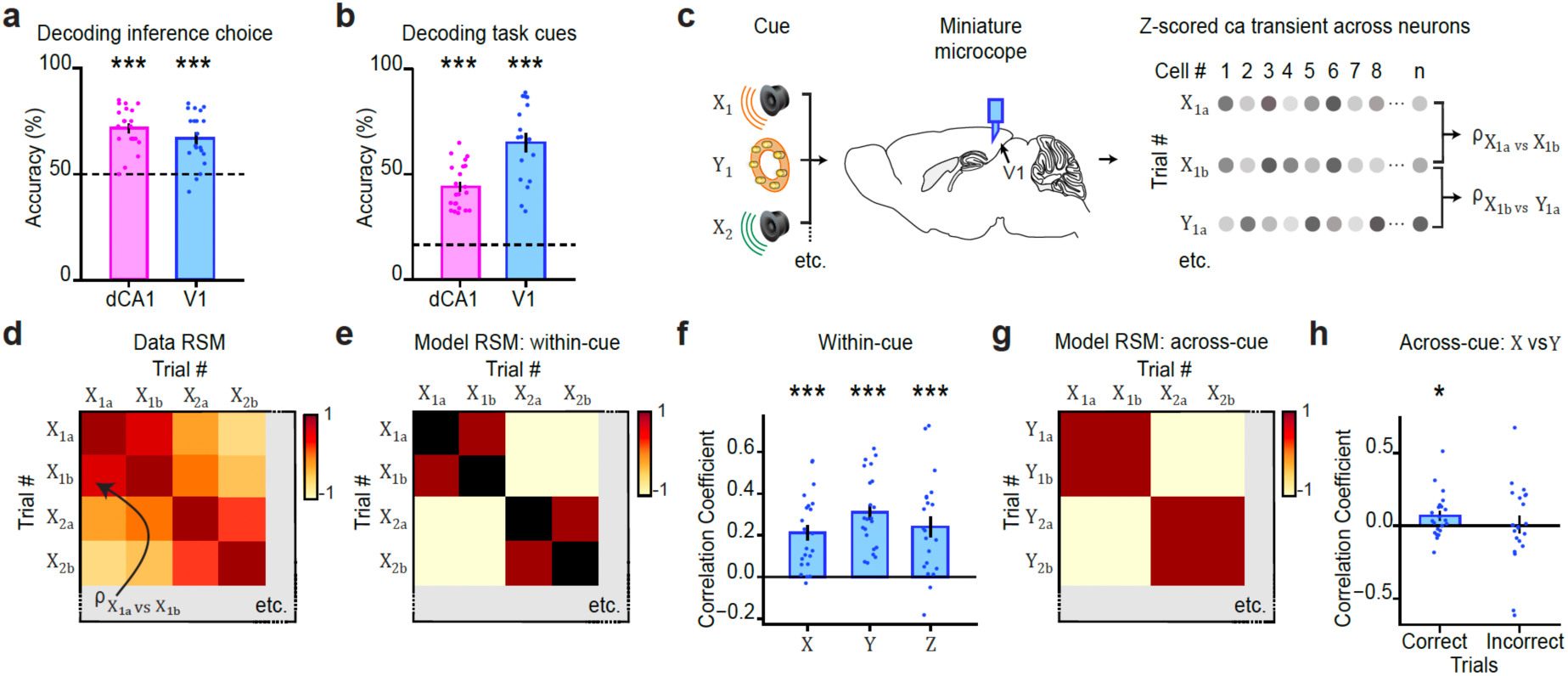
dCA1 and V1 represent task cues and engage a prospective code during inferential choice. **(a-b)** Cross-validated classifier with decoding accuracy significantly above chance for inference choice (*a,* dCA1: p<0.001; V1: p<0.001) and for task cue identity (*b,* dCA1: p<0.001; V1: p<0.001) when using population level data (left, pink: dCA1 spiking data from electrophysiology recordings; right, blue: V1 calcium transients from calcium-imaging data). Each point represents the mean classifier accuracy for one recording day. Bars, mean; error bars, ± SEM; dashed line, chance performance. **(c-e)** Schematics to illustrate RSA. **(c)** For each presentation of a cue (i.e. trial), a population vector was constructed from the average z-scored calcium transients recorded using calcium imaging in V1. The correlation coefficient between each pair of population vectors (i.e. for a pair of trials) was estimated. **(d)** The correlation coefficients were entered into a Data Representational Similarity Matrix (RSM). **(e-h)** Each Data RSM (*d*) was correlated with a Model RSM which defined the hypothesised information represented by population activity in V1. Equivalent analyses in dCA1 have been published previously^22^. **(e)** To test evidence for representation of the three difference cue types (auditory: *X_n;_* visual: *Y_n;_ outcome*: *Z_n)_* in V1, a “*within-cue*” discrimination model RSM was constructed to include positive correlation coefficients for cues within-set and negative correlation coefficients for cues across-set. A “*within-cue*” discrimination model RSM was constructed for each of the cue types, with an example shown here for auditory cues (*X_n)_* only. **(f)** The correlation between the Data RSM (e.g. *d*) constructed from calcium-imaging data in V1 and the Model RSM for “*within-cue*” discrimination (e.g. *e*) revealed significant discrimination of each type of task cue (auditory cues (*X_n)_*: p<0.001; visual cues (*Y_n)_*: p<0.001; outcome cues (*Z_n)_*: p<0.001). **(g)** To test evidence for a prospective code in V1, with predictive reinstatement of intermediary visual cues (*Y_n)_* in response to auditory cues (*X_n)_* in the inference test, an *“across-cue”* model RSM was constructed to include positive correlation coefficients for auditory (*X_n)_* and visual cues (*Y_n)_* within-set, and negative correlation coefficients for auditory (*X_n)_* and visual cues (*Y_1)_* across-set. **(h)** The correlation between the Data RSM constructed from calcium imaging data in V1 and the Model RSM for “across-cue” discrimination (*g*) revealed significant evidence for a prospective code, with predictive reinstatement of visual cues (*Y_n)_* in response to auditory cues (*X_n)_* in the inference test, on correct but not incorrect trials (correct: p=0.034; incorrect: p=0.868). **(f,h)** Each data point represents the correlation coefficient between the Data and Model RSMs for one recording day. Bars, mean; error bars, ± SEM. All p values generated using permutation testing.

**Supplementary Figure 2.**
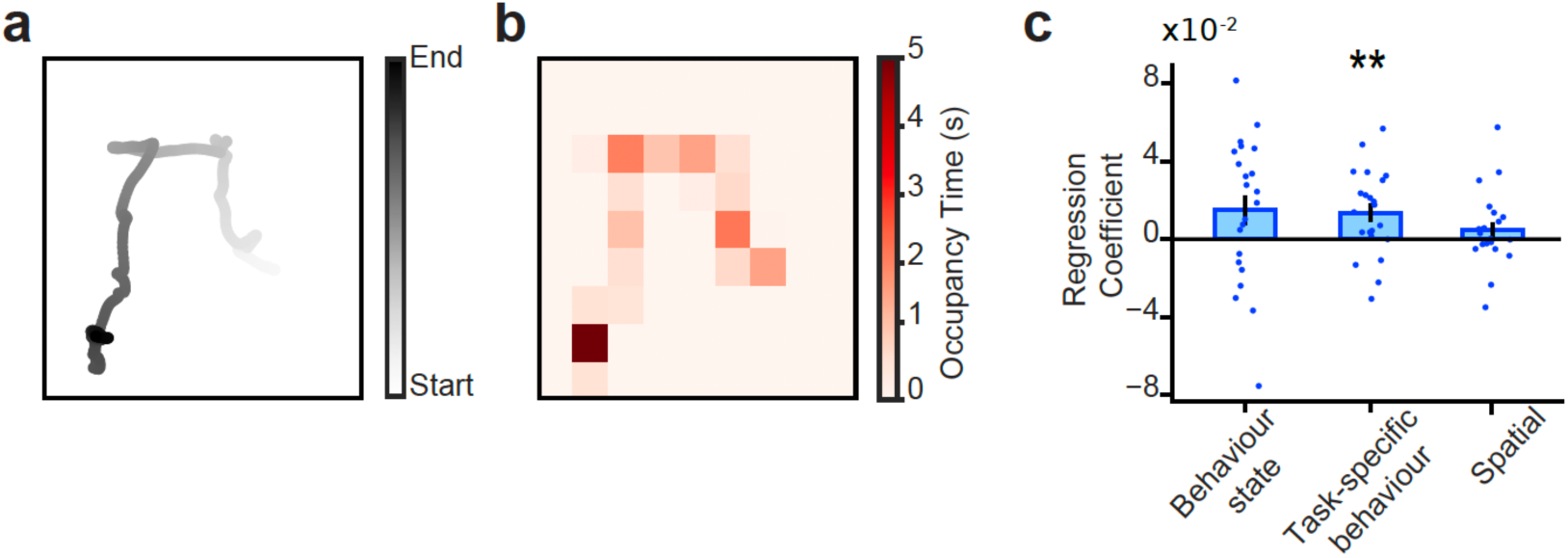
Spatial and behavioural controls for the RSA analysis shown in Figure 3. **(a)** Example trajectory of the mouse during a single presentation of a cue. **(b)** Occupancy map from the trajectory shown in *a* when split into 5cm^2^ bins. **(c)** To control for potential confounding effects of spatial representation in V1^52^, during the inference test we repeated the RSA analysis designed to test evidence for predictive reinstatement of visual cues (Y*_2)_* in response to auditory cues (X*_2)_* on correct but not incorrect trials (Figure 3g), now using linear regression to predict the RSM using three different possible explanatory model RSMs. For each recording day, we first generated a Data RSM (e.g. Figure 3c-d), before constructing three explanatory Model RSMs (e.g. Figure 3g), which included: *(1)* “Behaviour state”, a model designed to control for representational overlap in spatial trajectories leading to the dispenser. For this Model RSM, pairs of trials were highly correlated if the mouse made spatial trajectories towards the dispenser on both trials in the pair, while all other pairs of trials had zero correlation. *(2)* “Task-specific behaviour”, a model designed to reflect the task-specific associations in combination with the behaviour of the mouse. Therefore, pairs of trials from the same set had positive correlations (i.e. visual cues Y*_2_* and auditory cues X*_2)_* while pairs of trials from different sets had negative correlation (i.e. visual cues Y_$_and auditory cues X*_2)_*. Additionally, pairs of trials where at least one trial was incorrect had zero correlation. *(3)* “Spatial”, a model designed to control for the spatial trajectory of the animal. The model included the z-scored Pearson correlation coefficients between the spatial trajectories of the mouse across each pair of trials, where spatial trajectories for each trial were estimated using the occupancy within each 5cm x 5cm bin of the environment (as shown in *b*). These three explanatory Model RSMs (*1-3*) were regressed against the data RSM from each recording day. The beta coefficients for each of the three model RSMs are shown in *c*, where each data point represents the beta coefficient for a given recording day. The Data RSMs were significantly predicted by the “Task-specific behaviour” Model RSM (p=0.008, permutation test), with a trend observed for the “Behaviour state” Model RSM (p=0.075, permutation test), and no significant relationship observed for the “Spatial” Model RSM (p=0.278, permutation test). This suggests that in response to auditory cues (X*_2)_* in the inference test, V1 significantly reinstates the predicted visual cues (Y*_2)_* on correct trials, as opposed to the mere spatial trajectory of the animal. Bars: mean across days. Error bars: ± SEM.

Next, we sought to identify the content of information represented by dCA1 and V1 during the inference task. Specifically, we asked whether neural activity in dCA1 and V1 can represent the six task cues (set 1: *X_1,_ Y_1,_ Z_1;_* set 2: *X_2,_ Y_2,_ Z_2)_*. First, we trained a cross-validated logistic regression classifier to predict cue identity from neural activity recorded during visual cues (*Y_n)_* and outcome cues (*Z_n)_* in the conditioning trials, and during auditory cues (*X_n)_* in the inference test. In both dCA1 and V1, the identity of the presented cue could be significantly decoded (spiking activity in dCA1: mean accuracy 44.04%; calcium activity in V1: mean accuracy 65.05%) (Figure 3b). To further verify that V1 is able to separately discriminate auditory, visual and outcome cues across set 1 and 2, we used Representational Similarity Analysis (RSA)^44^ (Figure 3c-f). RSA quantifies the correlation coefficient between neuronal activity patterns (or population vectors) for each possible pair of cues. As expected for a visual cortical region, when applying RSA to V1 we observed significant evidence for between set discrimination of visual cues (*Y_1_* vs. *Y_2)_*. Importantly, RSA in V1 also revealed significant evidence for between set discrimination of auditory (*X_1_* vs. *X_2)_* and outcome cues (*Z_1_* vs. *Z_2)_* (Figure 3f). Thus, our data provide evidence to suggest that V1 neuronal activity can discriminate both visual and non-visual task variables^45^, consistent with a role for V1 in sensory processing that extends beyond visual processing^46,47^.

Having established that V1 can discriminate the 6 task cues, we next investigated whether V1 performs a computation to serve inference. During the inference test, previous data recorded from dCA1 in mice suggests the hippocampus prospectively represents visual cues *Y_n_* in response to auditory cues *X_n,_* thereby chaining *X_n_* to *Z_n22_*. This predictive activity is reminiscent of hippocampal spiking activity in the spatial domain, where activity can “sweep” ahead of an animal’s location and predict subsequent behaviour^23,48–51^. This predictive activity suggests the hippocampus is situated high within a hierarchical *generative* model capable of predicting ongoing sensory experience before sensorimotor feedback is available. If at the time of inference V1 inherits a prospective neural code from the hippocampus, predictive activity may be observed in V1 with reinstatement of the predicted visual cues (*Y_n)_* in response to auditory cues (*X_n)_*.

To test whether V1 reinstates the predicted visual cues (*Y_n)_*, we assessed whether the intermediary visual cue (*Y_n)_* can be decoded from neuronal activity in V1, during presentation of the auditory cues (*X_n)_* in the inference test. Using RSA, we compared the population activity from V1 during auditory cues (*X_n)_* in the inference test with activity from V1 during visual cues (*Y_n)_* in the conditioning. In the inference test, we found that when the correct outcome is inferred, as indicated by behaviour, neural activity during the auditory cue (*X_n)_* shows higher representational similarity with the associated visual cue (*Y_n)_*, compared to the non-associated (cross-set) visual cue (*Y*_$_) (Figure 3g-h). Therefore, during correct inference, neuronal activity in V1 reinstates the set-specific intermediary visual cue (*Y_n)_*. This reinstatement of the associated visual cue (*Y_n)_* was not observed on trials when the incorrect outcome was inferred (Figure 3h). Moreover, significant reinstatement of the associated visual cue (*Y_n)_* was observed during correctly inferred trials when controlling for the spatial trajectory taken by the animal, and mere occupancy at the outcome dispenser (Supplementary Figure 2). Therefore, at the time of inferential choice, exposure to auditory cues (*X_n)_* elicits a representation of the expected visual cues (*Y_n)_* in V1, analogous to the prospective code reported in hippocampus^22^. Together, hippocampus and V1 may therefore support inference by representing learned associations to ‘‘look ahead’’ and predict the short-term future.

### Coordinated activity across the hippocampus and neocortex during sleep

During online inferential choice, neural activity in V1 appears to mirror activity observed in the hippocampus^22^ by representing a prospective code. Next, we aimed to test whether neural codes in V1 may be inherited from hippocampus during offline periods of sleep. Hippocampal ripples provide a candidate mechanism to facilitate hippocampal-neocortical interactions during offline periods of sleep. Previous electrophysiology recordings on this inference task suggest that dCA1 spiking activity during ripples can co-activate task relevant cues (e.g. *X_1,_ Y_1,_ Z_1)_*, including co-activation of task relevant cues that represent the unobserved yet inferred relationship between *X_1_* and *Z_122_*. This dCA1 activity provides an example of *generative replay* where spiking activity during ripples deviates from direct experience by “joining the dots” between cues that represent an inferred relationship. In dCA1, this type of generative replay is only observed for inferred relationships that lead to reward, namely between *X_1_* and *Z_122_*.

Here, we asked whether generative replay in dCA1 can provide a training signal to task-relevant neocortical circuits, namely RSCg and V1. More specifically, we predicted that: (1) generative replay within dCA1 should propagate to V1; (2) generative replay observed in V1 should occur after that observed in dCA1; and (3) generative replay in V1 should become increasingly independent from dCA1 over time.

To test the first of these predictions, we investigated whether activity in task-relevant neocortical circuits (V1 and RSCg) is coordinated with dCA1 ripples. Using our two recording approaches (Figure 2f), we were able to detect ripples in dCA1 during post-task sleep (Figure 4a) while recording the responses of neuronal populations in RSCg and V1. We first assessed the response of ripple-modulated neocortical cells in the peri-ripple period (Figure 4b-c), before quantifying the proportion of cells showing significant modulation to dCA1 ripples (Figure 4d). In cells/units that were significantly ripple-modulated (see *Methods*), an increase in calcium transients was observed in V1 (Figure 4b) and an increase in spiking activity in RSCg and V1, with the peak neocortical spiking coinciding with the dCA1 ripple peak and notable ramping in spiking activity occurring prior to the dCA1 ripple peak (Figure 4c). In V1, we found that between 38.4% (estimated using calcium data) and 62.9% (estimated using spike data) of recorded cells were significantly modulated by ripples. We also calculated the proportion of significantly ripple-modulated cells in RSCg (51.2%) and dCA1 (84.4%) using our multi-site electrophysiology data. For all three brain regions (dCA1, RSCg, V1), significantly more cells were modulated by ripples than expected by chance (Figure 4d). Taken together, and consistent with previous evidence to suggest coordinated activity across the hippocampus and sensory neocortex during ripples^16–18^, our data demonstrates that task-relevant neocortical circuits are significantly modulated during dCA1 ripple events.

**Figure 4.**
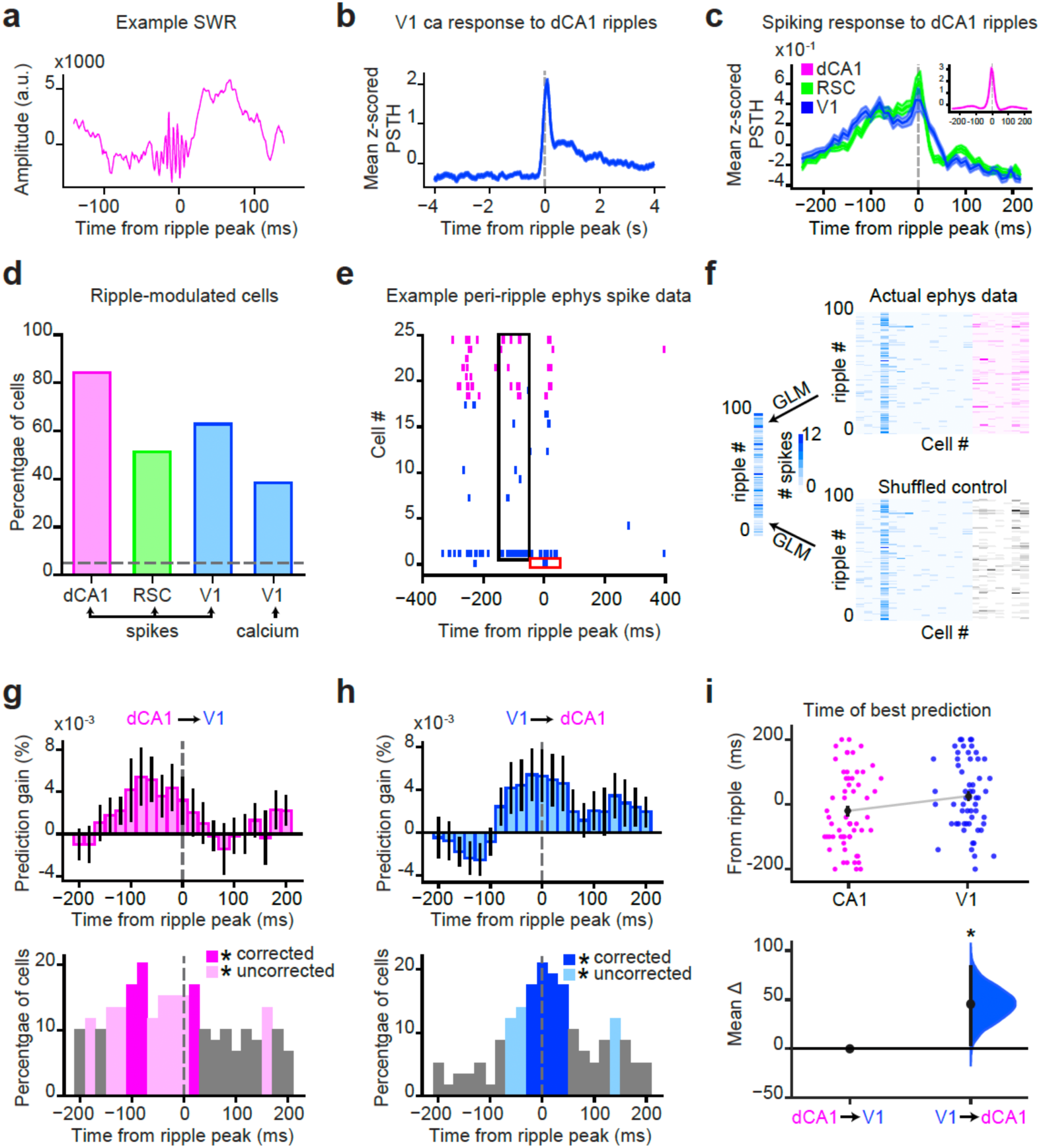
In ripples, hippocampal information content significantly predicts upcoming V1 information content. **(a)** An example SWR recorded from a tetrode in dCA1. **(b)** Average z-scored calcium transient of cells in V1 that are significantly modulated by ripples (see *Methods* for definition of “*ripple-modulated*”), in response to ripples recorded in dCA1. Mean ±SEM across cells. Dotted line, peak ripple power. **(c)** Average z-scored spiking response, recorded using electrophysiology, of cells/units that are significantly modulated by ripples in RSCg (green) and V1 (blue), in response to ripples recorded in dCA1. Inset shows the equivalent response recorded from cells/units in dCA1 (pink). Mean ±SEM across cells/units. Dotted line, peak ripple power. **(d)** Across all recorded cells/units, the total proportion of cells/units significantly modulated by ripples, for dCA1 (84.4%), RSCg (51.2%) and V1 (with separate proportions for spike-derived (62.9%) and ca-activity-derived (38.4%) cells). Dotted line, 0.05 chance threshold. **(e-f)** Schematic showing regression approach used for plots (*g-i)* to investigate whether in the peri-ripple period, information content in the hippocampus (“predictor”) can predict information content in neocortex (“target”) (and vice versa), using data recorded with electrophysiology. To this end, for each cell/unit in the “target” region, the spiking response is estimated across ripples, within a 100 ms sub-window centred on the peak ripple power. Next, we attempted to predict this spiking activity in the target region using a GLM. The GLM is designed to test whether activity in a “predictor” brain region can significantly predict the activity of the cell/unit in the “target” region, while controlling for activity from all other cells/units within the “target” region. The temporal dynamics of the relationship between the “predictor” and “target” brain regions are then assessed by constructing the GLM from different 100 ms data snapshots that span the 400 ms peri-ripple window centred on the peak ripple power. P-values are calculated by comparing the prediction scores against those obtained using a control that involves randomly shuffling the relationship between spikes and cell/units deriving from the “predictor” region, while retaining activity deriving from the other cells/units in the “target” region in a non-shuffled state (see *Methods*). **(g-h)** *Top:* Prediction gain, mean +/- SEM across all cells/units in the “target” region, recorded using electrophysiology. Spiking activity in the “predictor” region (*g*: dCA1; *h:* V1) is used to predict spiking activity in the “target” region (*g:* V1; *h:* dCA1), with spiking activity in the “predictor” region taken from shifted 100 ms sub-windows (each separated by 25 ms) that span the 400 ms peri-ripple window centred on the peak ripple power. Spiking activity in the “target” region is taken from the 100 ms sub-window centred on peak ripple power. Prediction gain indicates the difference in the prediction score between the actual data and the shuffled control (as illustrated in *f*). *Bottom:* For each shifted 100 ms sub-window (separated by 25 ms) used to estimate the response in the “predictor” region (*g*: dCA1; *h:* V1), the proportion of significantly predicted cells (p<0.05) in the “target” region (*g:* V1; *h:* dCA1) are indicated relative to a shuffled control. Dark coloured bars: significant relative to a Bonferroni corrected threshold of alpha=0.05/21 (21 = the number of shift bins). Light coloured bars: significant relative to a non-corrected threshold of alpha = 0.05. **(i)** *Top:* For each cell/unit in the “target” region (Left: V1; Right: dCA1), we estimated the time shift at which activity from the “predictor” region (Left: dCA1; Right: V1) gives the greatest difference between the actual and shuffled (control) GLM predictions. Black dots and bars: mean +/- STD. *Bottom:* A significant difference in means was observed for the optimal time shift for dCA1 predicting V1 (dCA1->V1, *left*) versus the optimal time shift for the reverse relationship, namely V1 predicting dCA1 (V1->dCA1, *right*) (permutation test: p=0.039), illustrated using bootstrap-coupled estimation (DABEST)^37^. This demonstrates an asymmetric sharing of information across the circuit where dCA1 significantly predicts upcoming V1 content during ripples. Effect size for the difference in means computed from 10,000 bias-corrected bootstrapped resamples^38^. Black dot, mean; black ticks, 95% confidence interval; filled-curve, sampling-error distribution.

### Hippocampal activity predicts upcoming cortical activity during ripples

During ripples recorded in hippocampus, we observed increased activity in task-relevant neocortical circuits (Figure 4b-c). We hypothesised that this increase in neocortical activity during ripples may reflect coordinated activity across hippocampal-neocortical circuits, where generative replay in the hippocampus has capacity to build a generative model across the cortical hierarchy. To test this hypothesis, we first asked whether the information content in dCA1 during ripples predicts subsequent information content in V1. Previous work has suggested that pre-ripple activity in sensory neocortex may predict upcoming dCA1 activity^17^. Indeed, in our electrophysiological recordings, we observed an increase in average spiking activity in V1 and RSCg prior to the peak of the dCA1 ripple (Figure 4c). However, if dCA1 provides a generative training signal to V1 during sleep, we would expect the *content* of spiking activity in dCA1 to predict upcoming activity in V1, rather than the reverse relationship.

To assess whether dCA1 ripple activity predicts upcoming V1 activity, or vice versa, we sought to capture the temporal dynamics of shared information content across hippocampal-neocortical circuits. To this end, we applied cross-validated general linear models to our multi-site electrophysiology data (Figure 4e-f). This involved training a model with the activity across cells/units in a given brain region (dCA1 or V1) during ripples, before assessing whether this model could predict the response of each cell/unit in a “target” region (V1 or dCA1, respectively). Critically, we controlled for activity across all other cells/units recorded in the target brain region to ensure any prediction from the model was specific to the precise content of spiking activity in the target region, rather than average changes in spiking activity. We then used a sliding window analysis to test at what jme and in what direction predictive activity occurred across the hippocampal-neocortical circuit, relative to peak ripple power recorded in dCA1. To summarise, this approach quantified how well we could predict the response of a given target cell/unit at the peak ripple power, using activity from another brain region that was shifted in time relative to the peak ripple power (Figure 4e-f).

To quantify whether predictions across the hippocampal-neocortical circuit were significant, we compared the results to a control distribution generated by shuffling spiking data in a manner which preserved the individual firing rates of any given cell, as well as the total activity across all neurons in response to a ripple (*see Methods*). Using this method, we assessed evidence for significant predictive information content across dCA1 and V1 (Figure 4g-i).

Firstly, for both pairs of cross-regional predictions (dCA1->V1, V1->dCA1), we observed more than one time window where the prediction was significant, even when controlling for multiple comparisons across time. This suggests that, during ripples, there is shared spiking content across task relevant hippocampal-neocortical circuits. When predicting V1 activity at the peak ripple power using dCA1 population activity (dCA1--> V1), we found that, on average, the best significant prediction occurred using dCA1 activity centred on a window 80ms before the ripple peak (Figure 4g). By contrast, for the reverse directionality (V1--> dCA1, Figure 4h), the best significant prediction occurred when V1 activity was centred on peak ripple power. This suggests an asymmetry in the direction of information flow in the peri-ripple period, whereby information is first represented in dCA1 before being represented in V1. Indeed, the optimal time shift for dCA1 predicting activity content in V1 occurred significantly earlier than the optimal time shift for the reverse relationship, where V1 predicts activity content in dCA1 (Figure 4i). This suggests that although *average* V1 activity increases prior to the dCA1 ripple (Figure 4c), activity content in V1 best predicts activity content in dCA1 at the peak ripple power (Figure 4h). By contrast, pre-ripple information content in dCA1 does significantly predict the content of V1 during the peak ripple power, over and above the reverse relationship (Figure 4g-i). This finding supports the idea that during ripples dCA1 activity provides a training signal to neocortex, to share information content across the cortical hierarchy.

### Memory reactivation in hippocampus predicts task-relevant reactivation in V1 during ripples

Next, we asked whether the shared information content across dCA1 and V1 during hippocampal ripples reflects processing of task-relevant information. During hippocampal ripples recorded during post-task sleep, we assessed the response of “cue cells”, namely those V1 cells identified using a GLM as responsive to task cues (Figure 5a). “Cue” cells in V1 (Figure 5b-c) were significantly more active during ripples compared to all other cells (Figure 5d). Therefore, V1 neurons that represent task cues online were preferentially active during offline ripples. In addition, building on findings reported in Figure 4, activity in hippocampal “cue” cells significantly predicted activity in neocortical “cue” cells across ripples (Figure 5e). These findings demonstrate coordinated task-relevant memory reactivation across dCA1 and V1 during ripples.

**Figure 5.**
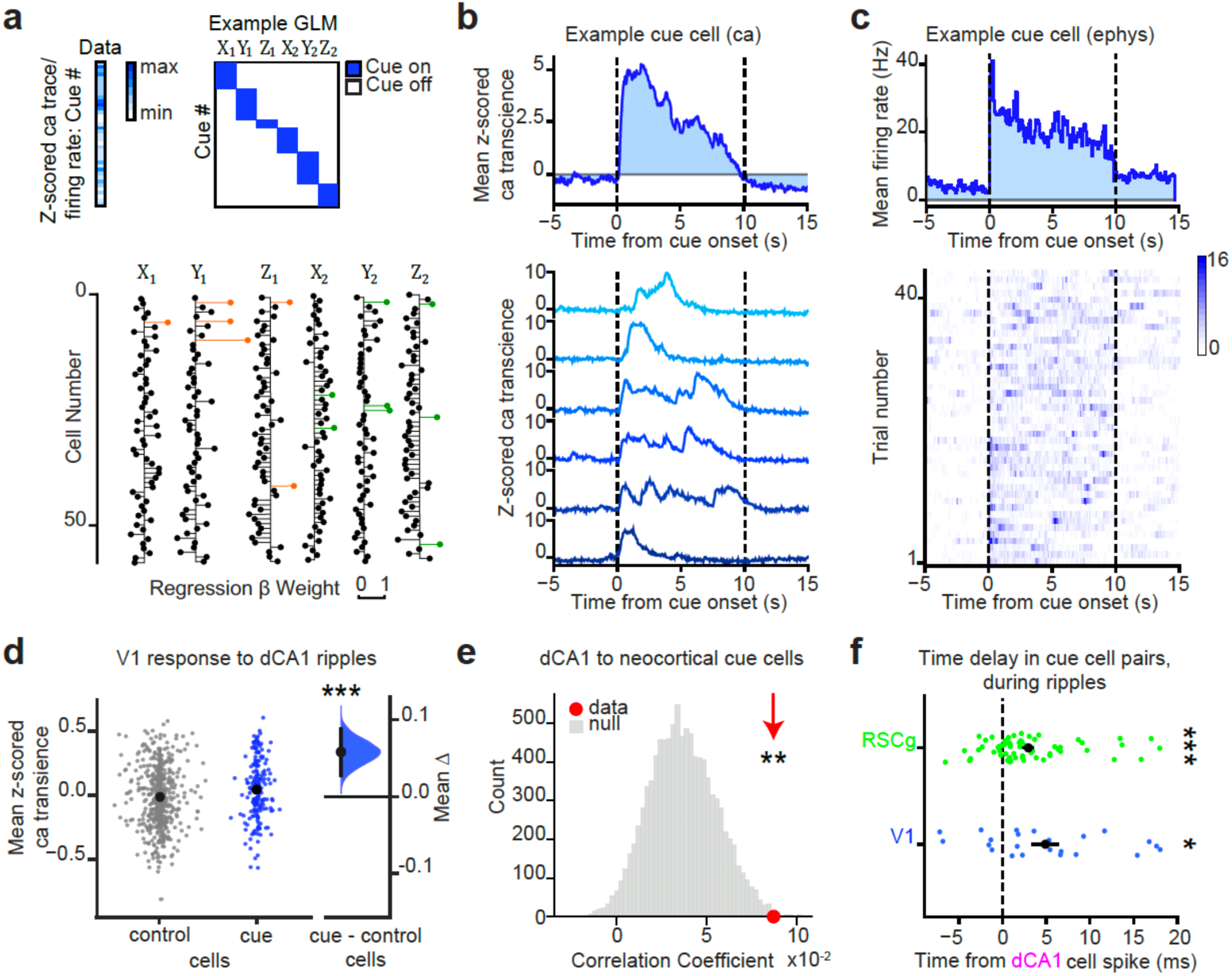
Cue cells are preferentially reactivated during ripples. **(a)** *Top:* Schematic illustrating how “cue” cells are defined. For each cell/unit, the average z-scored calcium transient (calcium imaging data) or firing rate (electrophysiology data) was regressed against a GLM with six regressors, one for each task cue. For each cell/unit, this generates a list of parameter estimates for each of the six cues, where each parameter estimate indicates the extent to which the activity of the cell/unit is modulated by presentation of the cue*. Bottom:* Example parameter estimates taken from the regression shown above, across all cells/units for the six task cues. Cells/units with parameter estimates greater than 2 standard deviations above the mean are highlighted in orange (set 1 cues) and green (set 2 cues). *Schematic adapted from*^22^. **(b-c)** Example “cue” cells from V1 calcium data (***b***, example V1 neuron in response to cue *Y_1)_* and from V1 electrophysiology data (***c***, example V1 neuron in response to cue *Y_2)_*. *Top:* mean response of the example “cue” cell to all presentations of the cue. *Bottom:* Activity of the “cue” cell in response to each individual presentation of the cue, showing z-scored calcium transient across six example trials (***b***), and number of spikes per time bin across all recorded presentations of the cue (***c***). Dashed vertical lines indicate the cue onset and offset. **(d)** Comparison of the response of V1 “cue” cells to all other “control” cells, during ripples detected in dCA1. Each data point represents the mean z-scored calcium transient for one cell/unit within 100 ms of peak ripple power. Difference in mean response to ripples for “cue” versus “control” cells is shown using a bootstrap-coupled estimation (DABEST) plot^37^. Effect size for the difference is computed from 10,000 bias-corrected bootstrapped resamples^38^. Black dot, mean; black ticks, 95% confidence interval; filled-curve, sampling-error distribution. “Cue” cells were significantly more responsive to ripples than “control” cells (permutation test: p<0.001). **(e)** Across ripples, the activity in dCA1 “cue” cells was significantly correlated with activity in neocortical (RSCg and V1) “cue” cells, relative to neocortical “control” cells. Null distribution shown in grey, generated by randomly selecting neocortical “control” cells and correlating their activity across ripples with the activity of “cue” cells in dCA1. Red dot and arrow indicate the correlation coefficient between neocortical “cue” cells and dCA1 “cue” cells, which were significantly more correlated than neocortical “control” and dCA1 “cue” (dCA1) cells (permutation test: p=0.007). **(f)** Time delay between spikes in pairs of “cue” cells during ripples, where one cell/unit in the pair derives from dCA1. For any given pair of “cue” cells: the first spike in dCA1 within a 400 ms time window centred on the peak ripple power was identified; similarly, the first spike in RSCg (top, green) and V1 (bottom, blue) were identified within this time window; across all ripples, the mean time difference between the first dCA1 spike and the first RSCg or V1 spike was estimated. On average, in the peri-ripple window, “cue” cells in both RSCg and V1 were significantly more likely to spike after “cue” cells in dCA1 (binomial tests: dCA1 vs V1: p=0.035, mean difference: 4.90 ms; dCA1 vs RSCg: p<0.001, mean difference: 2.95 ms). Each coloured dot represents the mean time difference for a unique cell pair. Black dots indicate the mean +/- SEM for each brain region.

Given evidence for shared task-related activity across hippocampus and V1 during ripples, we aimed to further characterise the temporal relationship between dCA1 and V1. We reasoned that if dCA1 provides a generative training signal to V1 during ripples, then task-relevant spiking activity in dCA1 should precede that in V1, thereby building on results shown in Figure 4g-i. To test this prediction, we compared the timing of spiking activity in “cue” cells identified in dCA1, RSCg and V1 using electrophysiology. We found that “cue” cells in dCA1 fired significantly earlier than “cue” cells in both RSCg (p<0.001) and V1 (p=0.030) (Figure 5f). This suggests that during ripples, task relevant memory reactivation occurs first in hippocampus before propagating to relevant sensory neocortical regions, namely RSCg and V1 in this case.

### Task-relevant sequential structure in V1

Having demonstrated that task-relevant memory reinstatement in dCA1 predicts corresponding reinstatement in V1 during ripples, next we asked whether this V1 memory reinstatement contains sequential structure. Specifically, we hypothesised that during periods of sleep, task cues would be reinstated in V1 *in sequence* to effectively chain together X*_n_* and Z*_n,_*despite these two cues never being experienced together in the awake state. Moreover, we predicted that this sequential structure in V1 should preferentially occur during ripples, when generative replay is otherwise observed in dCA1^22^.

To test this hypothesis we first used temporally delayed linear modelling (TDLM)^53^ to characterise sequences of internally generated neural representations in V1 during periods of sleep. TDLM is a domain general tool that uses a GLM framework to test evidence for sequential structure in neural data at the population level. Here, we apply the method to calcium imaging data acquired in V1, to measure evidence for sequential structure during sleep. To this end, we applied a cross-validated logistic regression classifier trained on ca transients recorded in response to the six task cues (*X_1,_ X_2,_ Y_1,_ Y_2,_ Z_1,_ Z_2)_* (Figure 6a-c), before applying the classifier to data acquired in the post-task sleep sessions. This generated a time course across the sleep session that indicated the reactivation probabilities for each of the six possible task cues (Figure 6d). We hypothesised that if V1 engages in generative replay, the decoded time courses should be structured such that reactivation of cue X*_n_* is chained to reactivation of cue Z*_n_* (*X_n_* → *Y_n_* → *Z_n)_*, despite these two cues never being experienced together during the task. Moreover, TDLM provides an opportunity to identify the time delay between sequential reactivation of each task cue. Thus, TDLM first estimates how likely a given cue is to be succeeded by reactivation of another cue at a lag of Δt (Figure 6e). Then, for each time lag Δt, TDLM estimates the extent to which the neural data follows a pre-specified set of transitions (e.g. *X_n_* → *Y_n,_ Y_n_* → *Z_n,_ X_n_* → *Z_n)_* otherwise termed “sequenceness” (Figure 6f). Thus, by applying TDLM to the decoded time courses, we could test evidence for structured sequences in the data recorded during sleep.

**Figure 6.**
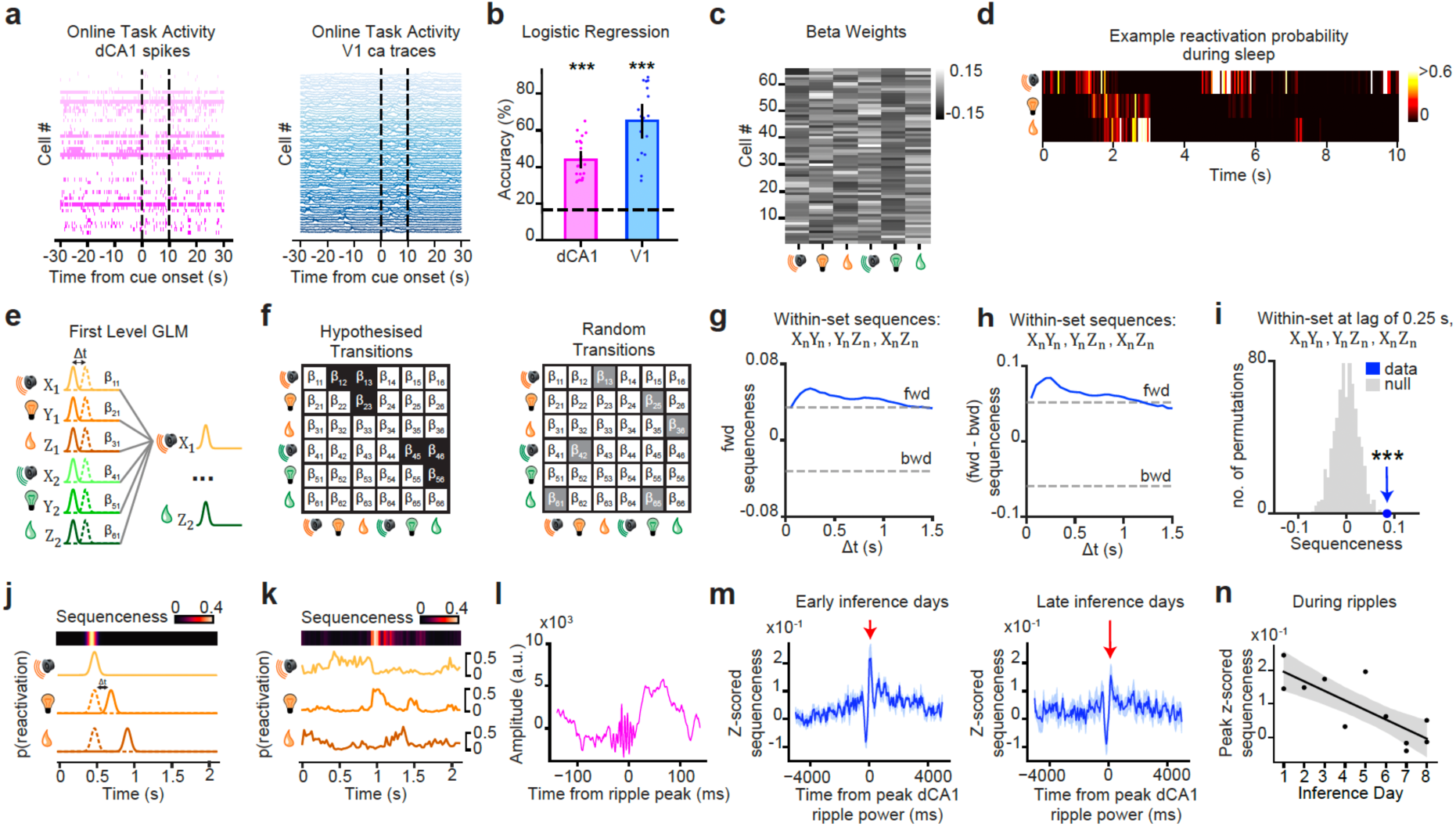
Measuring sequential replay during sleep. **(a)** Example spike trains from dCA1 (*left*, pink) or calcium traces from V1 (*right*, blue) during the task were used to train a cross-validated logistic regression decoder to predict the identity of the cue presented. **(b)** Cross-validated classifier decoding accuracy was significantly above chance for task cues, using spiking data from dCA1 (pink) and calcium data in V1 (blue) (dCA1: p<0.001; V1: p<0.001). Raw data points indicate average classifier accuracy for one recording day. Bars, mean; error bars, ± SEM; dashed line, chance performance. Note: this figure panel is a replica of Figure 3b. **(c)** Example beta weights from the logistic regression. **(d-n)** Temporal Difference Linear Modelling (TDLM)^53^ was used to quantify evidence for sequences in V1 calcium transients. **(d)** Beta weights from the logistic regression (*c*) were applied to sleep data to generate a time course of reactivation probabilities for each cue. An example snippet of these reactivation probabilities during sleep is shown for the putative reactivation of sequence (*X_n_* → *Y_n_* → *Z_n)_* from set 1. **(e)** Schematic showing how TDLM uses the time-lagged versions of the reactivation time courses from *d* to predict the reactivation time course of each other cue, using linear regression. This gives an empirical transition matrix at each time lag. **(f)** The empirical transition matrix is compared to a hypothesised transition matrix (*left*) based on the hypothesised set of task-related sequences (*X_n_* → *Y_n,_ Y_n_* → *Z_n,_ X_n_* → *Z_n)_*, to obtain a measure of “sequenceness”. A null distribution is generated by comparing the empirical transition matrix to 10,000 random permutations of the hypothesised transition matrix (*right*). **(g-h)** Evidence in V1 calcium transients for the hypothesised set of sequences (*X_n_* → *Y_n,_ Y_n_* → *Z_n,_ X_n_* → *Z_n,_* see *f)* across time lags. To control for multiple comparisons across each time lag, the statistical significance threshold, indicated by the grey dotted line, is set as the 2.5^th^/97.5^th^ percentile of all shuffles in the null distribution *(f)*, at the minimum/maximum value across all time lags. **(g)** Evidence for the hypothesised sequences *(f)* in the forward (*fwd*) direction, with a significant peak relative to the control distribution *(f)* at a time lag of 0.25 s (permutation test: p<0.001). **(h)** To control for autocorrelations, evidence for the hypothesised sequences *(f)* in the forward (*fwd*) direction was contrasted against evidence in the backwards (*bwd*) direction, giving the difference in sequenceness for the fwd-bwd direction with a significant peak relative to the control distribution *(f)* at a time lag of 0.25 s (permutation test: p<0.001). **(i)** Histogram showing significant evidence in V1 calcium transients for the difference in hypothesised sequences (*X_n_* → *Y_n,_ Y_n_* → *Z_n,_ X_n_* → *Z_n,_* see *f)* for the fwd-bwd direction (*h*) at the peak time lag of 0.25 s (data: blue), relative to the null distribution (null: grey) (permutation test: p<0.001). **(j)** Schematic showing that throughout the sleep session, TDLM can be used to obtain a timecourse of “sequenceness” using the optimal time lag identified in *g-h* (i.e. 0.25 s). **(k)** Example triplet sequence (*X_n_* → *Y_n_* → *Z_n)_* observed in V1 ca traces in sleep, together with “sequenceness” metric. **(l)** An example SWR recorded from a tetrode in dCA1, as shown in Figure 4a. **(m)** Average “sequenceness” in V1 ca traces, centred on peak ripple power. Dark blue: mean; Light blue: +/- SEM, across ripples. Left: early inference days (1-3); Right: late inference days (6-8). **(n)** Significant negative correlation between the peak z-scored V1 “sequenceness” during dCA1 ripples and the inference recording day (r=-0.79, p=0.004), where the “sequenceness” metric in V1 was referenced relative to a control distribution (see *Methods*). Note, across the entire sleep session, no significant change in average “sequenceness” was observed across recording days (r=-0.51, p=0.110). Black dots: each recording day; black line: linear regression model fit; shaded area: 95% confidence interval.

Importantly, by implementing a GLM-based approach, TDLM naturally allows for linear contrast between sequences that occur in the forward and backward direction (relative to the learned task structure). This provides an opportunity to control for potential spurious sequences attributed to representational overlap between task cues, which will equally affect both forward and backward sequences. Computing the difference in evidence for forwards and backwards “sequenceness” allows us to remove these autocorrelations.

Using TDLM, we found significant evidence for forward “sequenceness” in V1 calcium imaging data, relative to a null distribution generated by shuffling the hypothesised transition matrix (Figure 6f-g). This evidence for forward “sequenceness” remained significant when controlling for backward sequences (Figure 6h). The optimal time lag Δt for these sequential transitions occurred at 0.25 s, a plausible time lag given the relatively slow kinetics of the calcium indicator.

Having demonstrated evidence for sequential structure in V1 during sleep, next we asked whether this sequential structure preferentially occurs during dCA1 ripples. To test this, we quantified evidence for “sequenceness” in V1 calcium data across time, throughout the sleep session (Figure 6j-k), and assessed V1 “sequenceness” in the peri-ripple window. At the optimal time lag of 0.25 s we observed a peak in V1 “sequenceness” locked to the peak dCA1 ripple power (Figure 6l-m). However, the magnitude of the ripple-locked peak in V1 “sequenceness” significantly decreased across recording days (relative to a shuffled control), despite no significant change in overall V1 “sequenceness” across recording days (Figure 6m-n). These findings suggest that as task cues and the task structure become familiar, task-relevant sequential activity in V1 becomes increasingly decoupled from sequential activity in dCA1.

### Generative replay in V1 cue cells

Our analyses using TDLM provide evidence for sequential reactivation of task structure in V1. This sequential reactivation is locked to peak ripple power in dCA1, but with gradual decoupling over time. One limitation of TDLM is that it benefits from assessing evidence for a pre-specified *set* of hypothesised transitions. Thus, it was not possible to definitively establish whether cue X*_n_* is chained to reactivation of cue Z*_n,_* over and above replay of directly learned transitions. In a final set of analyses we tested evidence for specific sequences in V1, using calcium imaging data acquired from “cue” cells.

In both the spatial domain^23–27^, and during this inference task^22^, hippocampal activity during ripples shows sequential structure that extends beyond direct experience. Thus, hippocampal spiking activity can effectively “join-the-dots” to represent unobserved (yet logical) relationships between cues *X_n_* and *Z_n22_*. To test whether this hippocampal signature of generative replay propagates to V1, we calculated the probability that ripples nest calcium events from cue cells (Figure 7a) representing task cues (*X_n,_Y_n,_ Z_n)_*, including the unobserved yet inferred relationships (*X_n_* → *Z_n)_*.

**Figure 7.**
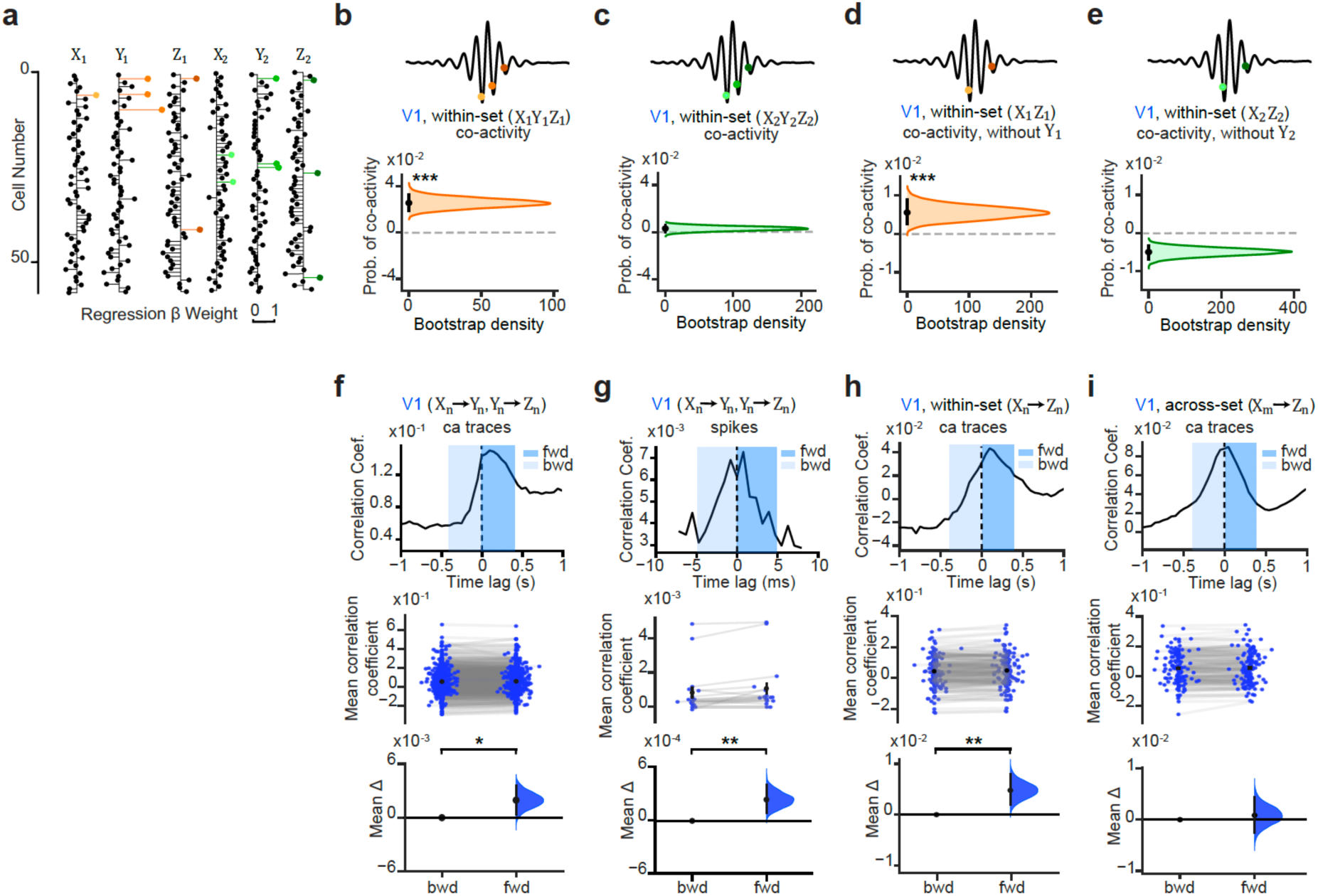
Generative sequences in V1 during sleep. **(a**) Schematic showing example cue cells for a given recording day, highlighted in orange (set 1 cues) and green (set 2 cues), where cue cells, as defined in Figure 5a and *Methods*. **(b-f)** Co-activity between triplets or doublets of cue cells in V1, during dCA1 ripples recorded during sleep. Co-activity was estimated using the probability of a calcium event in V1 occurring for a triplet or doublet of cue cells within the ripple window, where the ripple window was defined from electrophysiology data recorded in dCA1. The effect size for the difference between ripple and non-ripple windows is shown using a DABEST plot: black dot, mean; block ticks, 95% confidence interval; filled-curve, sampling-error distribution. **(b)** During dCA1 ripples, significant V1 co-activity was observed between triplets of cue cells from set 1 (*X_n_* & *Y_n_* & *Z_n)_*, namely the set that included reward (*Z_n)_* (permutation test, significantly more co-activity relative to null: p<0.001). **(c)** During dCA1 ripples, no significant V1 co-activity was observed between triplets of cue cells from set 2 (*X_2_* & *Y_2_* & *Z_2)_*, namely the set that included only a neutral outcome (*Z_2)_* (permutation test, significantly less co-activity relative to null: p<0.001). **(d)** During dCA1 ripples, significant V1 co-activity was observed between doublets of cue cells representing the inferred relationship from set 1 (*X_n_* & *Z_n)_* in the absence of *Y_n_* (permutation test, significantly more co-activity relative to null: p<0.001). **(e)** During dCA1 ripples, no significant V1 co-activity was observed between doublets of cue cells representing the inferred relationship from set 2 (*X_2_* & *Z_2)_* in the absence of *Y_2_* (permutation test, significantly less co-activity relative to null: p<0.001). **(f-i)** The temporal relationship between the z-scored activity of pairs of cues cells (as defined in Figure 5 and *Methods*) across the sleep session, quantified using cross-correlograms. In each case, the activity of the second cell in the pair was shifted while holding the activity of the first cell in the pair fixed, as reference, such that: positive time lags indicate *forward* (*fwd*) sequences; negative time lags indicate *backward* (*bwd*) sequences. *Upper:* Example cross-correlogram for a given pair of cue cells. *Middle*: mean correlation coefficients for *fwd* and *bwd* lags. Coloured dots: mean for a given pair of cue cells; black dot: mean; black tick: +/- SEM. *Lower:* Effect size for the difference in means for *fwd* versus *bwd*, computed from 10,000 bias-corrected bootstrapped resamples^38^: black dot, mean; black ticks, 95% confidence interval; filled-curve, sampling-error distribution. **(f-g)** In V1 ca-imaging data (*f*) and V1 electrophysiology data (*g*), we observed significant asymmetry in the cross-correlogram for within-set cue cell pairs representing learned associations (*X_n_* → *Y_n,_ Y_n_* → *Z_n)_*, indicating significant bias for *fwd* sequences (permutation tests, ca traces in *f*: p=0.028; spikes in *g*: p=0.008). **(h)** In V1 ca-imaging data, we observed significant asymmetry in the cross-correlogram for within-set cue cell pairs representing inferred relationships (*X_n_* → *Z_n)_*, indicating significant bias for *fwd* sequences (permutation test, ca traces: p=0.004). **(i)** In V1 ca-imaging data, we observed no significant evidence for asymmetry in the cross-correlogram for across-set cue cell pairs representing inferred relationships (*X_1_* → *Z_n)_* (permutation test, ca traces: p=0.655).

In V1, during dCA1 ripples recorded in sleep, we observed significant co-activity between cells representing all three cues from set 1 (*X_1_* & *Y_1_* & *Z_1)_* (Figure 7b), but not from set 2 (*X_2_* & *Y_2_* & *Z_2)_* (Figure 7C). The prioritisation of reward-related information is consistent with work investigating hippocampal replay of previous experience^54^, and with evidence that when mice are trained on this inference task, dCA1 prioritises sequences for the rewarded set^22^. Moreover, as observed in dCA1^22^, this result did not merely reflect simulation of the learned associative chain (*X_1_* → *Y_1_* → *Z_1)_*. Rather, in V1 we also observed significant co-activity between doublets of cue cells representing the inferred relationship from set 1 (*X_1_* & *Z_1)_* in the absence of the intermediary *Y_1_* (Figure 7d). Notably, no significant V1 co-activity was observed between cue cells representing the equivalent relationship for set 2 (*X_2_* & *Z_2)_* (Figure 7e). Taken together, these results suggest that during ripples, activity in V1 represents the unobserved yet inferred relationship that leads to reward, providing a signature of generative replay that mirrors that observed in dCA1.

To further verify that V1 shows significant evidence for sequences, we assessed the temporal relationships between activity in pairs of “cue” cells directly (Figure 7a) using cross-correlograms. Using both calcium imaging and electrophysiology data, we show that replay of learned associations during sleep is biased towards the forward direction in V1 (*X_n_* → *Y_n,_ Y_n_* → *Z_n;_* Figure 7f-g). Moreover, this bias towards forward sequences can also be observed for the inferred relationship (*X_n_* → *Z_n)_* (Figure 7h), but not for across-set relationships (Figure 7i). Together, these analyses provide evidence to suggest that replay in V1 preserves the learned associative task structure while also representing inferred relationships to provide a signature of generative replay.

## Discussion

Flexible behaviour depends on predictions that extend beyond direct experience, but the neural computations that generate these predictions have remained unknown. Here, we show that during sleep, neuronal sequences in V1 represent rewarding yet unobserved relationships that have not been directly experienced. Moreover, by simultaneously recording from hippocampus and V1, we show that information content in V1 during sleep is predicted by that in dCA1, but not vice versa. These findings provide evidence for “*generative replay*” outside the hippocampus and suggest that generative replay in dCA1 may propagate through neocortical circuits. Overall, our findings suggest that the sleeping brain has a remarkable ability to self-generate training data. This training data may then be used to construct a deep hierarchical generative model capable of formulating predictions that stretch beyond direct experience.

To test evidence for sequential activity in V1 we use several approaches. Previous studies in rodents have typically relied on decoding memory identity^55^, or have assessed evidence for sequences across sleep frames using multi-unit ensemble activity^56^. In humans, sequential replay has been detected in V1 using fMRI paoerns, at a relatively low jme resolution^57^. Here, in mouse V1 we provide the first evidence for *single-unit* sequential replay across cells in V1, and demonstrate that *genera7ve replay* can be observed in V1, where sequences go beyond those that have been directly observed to represent profitable (rewarding) inferred relationships (i.e. *X_1_* → *Z_1)_*. While our data suggest that generative replay is inherited from sequences in dCA1, over time we observe a gradual decoupling between V1 sequential activity and dCA1 ripples, suggesting that once V1 has inherited neural codes from dCA1, V1 itself may contain an internal model capable of synthesising novel sequences of predicted features. These findings align with predictive coding theory where V1 is proposed as an active participant in generative processing^58–60^.

Interestingly, both replay and generative replay sequences in V1 are biased towards the forward direction, relative to the learned temporal statistics of the task. These forward sequences may be necessary to establish a generative model capable of predicting *forthcoming* behaviour in a manner consistent with model-based sequential planning^61,62^, where future events can be anticipated even if they have not previously been sampled^25,63^. Intriguingly, the directionality of sequences in V1 contrasts with the dominant directionality in dCA1 where examples of generative replay on the same task are biased towards the reverse direction^22^. To reconcile these differences in replay directionality across dCA1 and V1, we suggest that a division of labour may occur across ripples, or even within a ripple. For example, during ripples where the content of dCA1 biases that in V1, dCA1 spiking sequences may be biased towards the forward direction. By contrast, during ripples where the content of dCA1 biases reward regions, dCA1 spiking sequences may be biased towards the backward direction, to allow credit-assignment^64^ where value is back-propagated to previously neutral cues^22^.

We carefully quantify the temporal relationship of shared information content across dCA1 and V1. Previous studies have shown evidence for coordinated reactivation of events across both hippocampus and V1^16,55^, including an increase in content coherence across these two brain regions in the pre-ripple window^55^. However, the temporal directionality of information flow between dCA1 and V1 during reactivation has not previously been established. Other reports have examined the temporal relationship between the hippocampus and primary auditory cortex^65^, but without controlling for general increases in multi-unit activity during ripples. Here, to determine the temporal directionality of precise information exchange between dCA1 and V1, we implemented a regression analysis that controls for: *(1)* the response of all other cells/units within the target region; *(2)* the population rate of the predictor region within each ripple, together with the firing rate of each neuron in this region. Overall, this approach controls for the increase in multi-unit activity during ripples, thereby ensuring that prediction scores are directional and not artificially inflated by elevated spike counts in the ripple window. Applying our approach to the peri-ripple time window, we show that while activity in dCA1 can predict subsequent activity in V1, we see no significant evidence for the reverse relationship. Therefore, despite average increases in V1 neural activity prior to the ripple, the precise information content in V1 during this period does not significantly predict upcoming information content in dCA1, over and above general increases in multi-unit activity. These findings are consistent with the view that dCA1 mediates the content of memory consolidation^1,2^, providing a putative index^66^ or training signal to sensory cortical brain.

To identify a candidate hippocampal-neocortical circuit that supports inference, we used anatomical tract tracing in mice. We show putative evidence for a disynaptic anatomical pathway from dCA1 to V1, via the RSCg. Notably, the same three brain regions selectively show a significant increase in activity for correct compared to incorrect inference when humans perform the same task^22^. This suggests that during correct inference, hippocampal activity is modulated together with brain regions important for memory and the processing of relevant sensory cues. Our anatomical findings further show that dCA1, RSCg and V1 form a putative disynaptic anatomical circuit, providing a pathway for hippocampal activity to influence activity in V1. This evidence for a disynaptic pathway builds on previous work in rodents that suggests evidence for a monosynaptic connection between dCA1 and RSCg^40,67^, and a monosynaptic connection between RSC and V1. Our anatomical characterisation of RSC further supports the view that RSC provides a prominent gateway for information exchange between the hippocampal formation and areas of neocortex.

Using both electrophysiology and calcium imaging to functionally characterise activity across the dCA1, RSCg and V1 circuit, we show that V1 is able to represent auditory and outcome cues, in addition to visual cues. V1 is one of the earliest brain regions in the cortical hierarchy. Historically, neurons in V1 have been recognised for their role in processing visual information, showing selectivity in orientation, spatial frequency, temporal frequency, and in the direction of visual motion^68–72^. However, several recent studies suggest V1 may encode richer information. For example, neurons in V1 encode both sensory and non-sensory information^45,47^ supporting learning^73^ and memory^46^. Moreover, visual responses in V1 are influenced by navigational signals, which are coherent with those encoded in hippocampus and reflect the animal’s subjective position^52,74^. Our findings support this broader characterisation of V1, implicating V1 in representing sensory cues that are not merely visual.

Taken together our findings suggest that during ripples in sleep, generative replay represented by dCA1 propagates to sensory cortical circuits, including V1. This allows V1 to represent profitable (rewarding) yet unobserved relationships that extend beyond experience. These findings suggest that a core feature of memory consolidation involves the fundamental reorganisation of information content across cortex, to build novel predictions that may be profitable in the future. Generative replay in hippocampus may therefore endow neocortex with the capacity to build a hierarchical generative model that supports our most flexible and adaptive behaviours.

## Methods

### Animals

16 adult mice were included in the study (age 4-6 months, males) that were heterozygous for the transgene expressing Cre recombinase, with expression driven by the CaMKIIa promoter maintained on a C57BL/6J background (RRID: IMSR_JAX:005359), or C57BL/6J (Charles River UK). Animals had free access to water in a dedicated housing facility with a 12/12h light/dark cycle (lights on at 07:00h). Animals were housed with their littermates up until the start of the experiment, during which they were housed alone. Food was available *ad libitum* prior to the start of the experiments. All experiments involving mice were conducted according to the UK Animals (Scientific Procedures) Act 1986 under personal and project licenses issued by the Home Office following ethical review.

### Surgical Procedures

Mice received viral injections and microdrive implantations under gaseous isoflurane anaesthesia (∼1% in 1 L/min O*_2)_*, with systemic and local analgesia administered subcutaneously (meloxicam 5 mg/kg; buprenorphine 0.1 mg/kg).

#### Viral injections

For all viral injections, we used a glass micropipette with a delivery rate of 100 nl/min in dCA1; 50 nl/min in RSCg; 50 nl/min in V1. Following virus delivery, the pipette was held in place for 5 mins before being withdrawn. Viral injections were targeted to bilateral dCA1 using stereotaxic coordinates (-1.7/-2.3 mm anteroposterior from bregma, ±1.7 mm lateral from bregma, and -1.1 mm ventral from the brain surface), to bilateral RSCg using stereotaxic coordinates (-1.7/-2.3 mm anteroposterior from bregma, ± 0.15 mm lateral from bregma, and –0.2/0.4 mm ventral from the brain surface), and to V1 using stereotaxic coordinates (+0.3/+0.7mm anteroposterior from lambda, +3.0 mm medial-lateral from lambda, and -0.4/-0.15 mm ventral from the brain surface). To visualise the pathway from dCA1 to RSCg, in CaMKIIa-Cre mice we injected a Cre-dependent tdTomato (rAAV2-CAG-Flex-tdTomato; UNC; AV4598c) into dCA1 (350 nl per site), and GFP (rAAV2-CAG-FLEX-ArchT-GFP; UNC; AV4984b) into RSCg (150 nl per site) of the same mouse, using viruses from E.S. Boyden (available at UNC Vector Core) (n=2, Figure 2b-c).

To visualise the putative di-synaptic pathway from dCA1 to V1, an intersectional trans-synaptic strategy was employed (n=3, Figure 2d-e), with details on cloning the plasmids, cell culture and transfection described previously^43^. The strategy involved a double-conditional Boolean logic scheme of transgene expression, where translation of the fDIO-GFP construct is conditional on the sequential activity of Cre and flippase (FlpO) recombinases within the same cell, meaning that GFP is only expressed in the presence of both viruses (Figure 2d). Therefore, the two recombinases act in series: the presence of Cre allows the expression of FlpO, which in turn allows the expression of GFP. We packaged plasmids containing the Cre-dependent FlpO transgene (cDIO-FlpO) in a high-titer serotype 1 AAV vector capable of anterograde trans-synaptic transfer (AAV1-EF1a-cDIO-FLPo-Myc; constructed in house: packaged in HEK293 cells with AAV1 coat proteins (1.6E13 vg/mL) using AAV-1 Helper Free Packaging System vectors VPK-401 vector kit, Cell Biolabs Inc., USA)^43^, which we injected into bilateral dCA1 of CamKII-Cre mice (250nl per site). The second AAV harbouring the Flpo-dependent GFP (AAV2-EF1a-fDIO-EGFP, packaged in UNC Vector Core, 9.6E12 vg/mL) was then injected in the RSCg of the same mice (150nl per depth per site). To control for non-specific expression due to pipette-leakage or spontaneous recombination, we repeated the procedure without injecting the first virus (AAV1-EF1a-cDIO-FLPo-Myc) into dCA1 (n=2, Supplementary Figure 1b-c), despite loading the pipette and lowering it to the target dCA1 coordinate (Supplementary Figure 1b).

#### Microdrive implantation

For triple (n=2) and dual (n=3) site electrophysiological recordings (Figure 2f, 2l), mice were implanted with a microdrive containing 12 independently moveable tetrodes that targeted the pyramidal cell layer of bilateral dCA1 in the hippocampus using stereotaxic coordinates (-2.0 mm anteroposterior from bregma, ±1.7 mm lateral from bregma, and -1.1 mm ventral from the brain surface), bilateral RSCg using stereotaxic coordinates (-2.0 mm anteroposterior from bregma, ± 0.15 mm lateral from bregma, and -0.4 mm ventral from the brain surface), and bilateral V1 using stereotaxic coordinates (+0.5 mm anteroposterior from lambda, ±3.0 mm lateral from lambda, and - 0.25 mm ventral from the brain surface). The minimum distance between neighbouring tetrodes inserted in a given brain region of each hemisphere was 0.4 mm. Tetrodes were constructed by twisting together four insulated tungsten wires (12.7 μm diameter, California Fine Wire, CA, USA) which were briefly heated to bind them together into a single bundle. Each tetrode was loaded in one cannula attached to an M1.0 screw to enable independent depth manipulation. The microdrive was implanted under stereotaxic control with reference to bregma and lambda, using the central target coordinates as a reference point to position each individual tetrode contained in the microdrive, with initial implantation immediately above the pyramidal cell layer (dCA1), or dorsal to the target coordinate (RSCg and V1). Following the implantation, the exposed parts of the tetrodes were covered with paraffin wax, after which the drive was secured to the skull using dental cement. For extra stability, 3-4 stainless-steel anchor screws were inserted into the skull before the drive was implanted. Two of the anchor screws, were attached to 50 μm tungsten wires (California Fine Wire) and further served as a ground and reference electrodes during the recordings.

For integrated calcium imaging and tetrode recordings (n = 4 mice, Figure 2f-h), the adeno-associated viral (AAV) vector GCaMP6f (CaMKII-GCaMP6f-WPRE-SV40; UPenn Vector Core) was injected in left V1 (150nl per depth per site). Two weeks after viral injection, mice were implanted with a microdrive containing 2-4 independently moveable tetrodes (as described above) that targeted the pyramidal layer of right CA1 in the hippocampus using stereotaxic coordinates (-2.0 mm anteroposterior from bregma, +1.7 mm lateral from bregma, and -1.1 mm ventral from the brain surface), and a prism lens (Inscopix ProView Prism Integrated Lens: 1.0 mm x 4.3 mm; product code: 1050-005301) targeting left V1 using stereotaxic coordinates (+0.5 mm anteroposterior from lambda, +3.0 mm lateral from lambda, and -0.5 mm ventral from the brain surface). Following the implantation of the integrated microdrive and prism lens, the drive was secured to the skull using dental cement and the baseplate to the integrated prism lens sealed with a baseplate cover (Inscopix).

### Tissue processing and immunohistochemistry

At the completion of experiments, all mice were anaesthetised with pentobarbital and transcardially perfused with 0.1 M PBS followed by 4% paraformaldehyde (PFA) in PBS, at 10 ml/min. Chronic implants were removed, if present. Brains were extracted and kept in 4% PFA for at least 24 hours and then transferred to PBS (with 0.05% sodium azide). For immunohistochemistry, free-floating sections (50 μm) were rinsed in PBS (3 x 10 mins) before being blocked for 1 hour at ∼20°C in 0.3% Triton X-100 (PBS-T) with 10% Normal Donkey Serum (NDS). Sections were then incubated for 48 hours at 4°C with primary antibodies diluted in PBS-T and 1% NDS blocking solution (GFP anti-chicken, 1:1,000, Aves Labs, catalogue no. GFP-1020; RFP anti-rat, 1:1,000, ChromoTek, catalogue no. 5f8; PV anti-guinea-pig, 1:2000, Synaptic Systems, catalogue no. 195004). Sections were then rinsed in PBS-T (3 x 10 mins) and incubated overnight at 4°C in PBS-T and 1% NDS blocking solution containing secondary antibodies (488 donkey anti-chicken, 1:1,000, Jackson ImmunoResearch, product ref: 703-545-155; Cy3 donkey anti-rat, 1:1,000, Jackson ImmunoResearch, product ref: 712-165-153; 405 donkey anti-guinea-pig, 1:100, Jackson ImmunoResearch, product ref: 706-475-148). After rinsing in PBS (3 x 10 mins), sections were counterstained for 10 mins with 0.01% DAPI in PBS-T, before being rinsed again in PBS (3 x 10 mins). Sections were mounted on non-gelatinised glass slides, cover-slipped with Vectashield mounting medium (Vector Laboratories, catalogue no. H-1000) and stored at 4°C. Images were acquired using a Zeiss confocal microscope (LSM 880) with a Plan-Apochromat x20/0.8 objective and the ZEN Black software (v 2.3) (Figure 2, Supplementary Figure 1).

### Behavioural Task

For the in vivo recordings (n=9 mice), the mice performed an inference task, as described previously^22^. The task consisted of 3 stages (Figure 1a): an observational learning stage (6 consecutive days), a conditioning stage (4 consecutive days), and an inference test stage (5-8 consecutive days) (Figure 1c). The mice were fed *ad libitum* until day 4 of the observational learning stage. At this point, the mice were food-deprived to 90% of their free-feeding body weight to prepare for the conditioning and inference test stages. The task was performed in a walled, open field enclosure (41 x 41 cm square box with 38 cm high walls) in which the mice were allowed to freely move around (Figure 1b). Presentation of the sensory cues was controlled by a single-board microcontroller (Arduino Mega 2560 Rev3). The auditory cues (*X_n)_* were delivered through two speakers placed above the open field. Two LED panels used to provide visual cues (*Y_n)_* were affixed on opposite walls of the open field. A liquid dispenser and aspirator fitted with an infrared beam for lick detection were used to deliver and remove the outcomes (*Z_n)_*. The dispenser was fixed to the wall between the LED panels so that the LEDs were approximately equidistant from the dispenser (Figure 1b). Across all stages of the task, trials were only triggered when the animal was moving and not immediately in front of the dispenser. Across training days, and across the inference test, cues from sets 1 and 2 were presented equally often.

During the observational learning stage, the mice were exposed to repeated pairings of auditory (*X_n)_* and visual (*Y_n)_* cues (Figure 1a). Each auditory cue lasted 10 s and was immediately followed by a visual cue which lasted 8 s. There were two auditory-visual associations (set 1: *X_1_* → *Y_1;_* set 2: *X_2_* → *Y_2)_* and the pairings between cues were randomly assigned for each mouse. The auditory cues were pure tones (10 kHz and 2 kHz) and the visual cues were sets of coloured LEDs (L-shaped green strip and circular orange set) (Figure 1b). Each day, the mice were placed in the open field arena for 8-10 training sessions, each lasting ∼20 mins. In each training session, 12 trials (*X_n_* → *Y_n)_* were delivered with ∼1.5 mins inter-trial interval. On training days 1-4, trials presented within session were blocked by set (*X_1_* → *Y_1_* or *X_2_* → *Y_2)_*. On the final two training days 5-6, trials presented within session were presented in a pseudo-random order across the two sets (*X_1_* → *Y_1_* and *X_2_* → *Y_2)_*.

During the conditioning stage, the animals were re-exposed to the visual cues (*Y_n)_* from the observational learning stage, which were now paired with an outcome (*Z_n)_* (Figure 1a). The outcome cues constituted either a drop of 15% sucrose solution (*Z_1,_* rewarding outcome), or a drop of water (*Z_2,_* neutral outcome). Each visual cue was presented for 8 s and was immediately followed by sucrose or water delivery at the outcome dispenser, made available for 10 seconds before being removed (Figure 1d). The conditioning stage began immediately following the end of the observational learning stage. Each day, the mice were placed in the open field arena for 9-10 training sessions, each lasting ∼20 mins. In each training session, 12 trials (*Y_n_* → *Z_n)_* were delivered with ∼1.5 mins inter-trial interval. On training day 7-8 (day 1-2 of conditioning stage), trials presented within session were blocked by set (*Y_1_* → *Z_1_* or *Y_2_* → *Z_2)_*. On training days 9-10 (days 3-4 or conditioning stage), trials presented within session were presented in pseudo-random order (*Y_1_* → *Z_1_* and *Y_2_* → *Z_2)_*. The two visual-outcome associations (set 1: *Y_1_* → *Z_1;_* set 2: *Y_2_* → *Z_2)_* and the pairings between cues were randomly assigned for each mouse.

In the inference test, the mice were exposed to the auditory cues (*X_n)_* in isolation for 10 s (Figure 1a, 1c). Based on learning from the previous days, mice could infer the expected outcome (*Z_n)_*. The outcome was never delivered in response to the auditory cue (*X_n)_*. The inference stage began immediately following the end of the conditioning stage. Each day, the mice were first reconditioned (set 1: *Y_1_* → *Z_1;_* set 2: *Y_2_* → *Z_2)_*, with exposure to two consecutive ∼20 min sessions, each with at least 12 trials presented in a pseudo-random order across set. Reconditioning sessions were followed by the inference test, where 8-12 trials were presented using a pseudo-random order across sets. At the end of the inference test, animals were removed from the open field to rest before returning for a brief session of reconditioning trials followed by the inference test. This sequence of reconditioning followed by inference test was then repeated a third time (Figure 1c). Within a given day, a total of 28 inference trials were presented. By interleaving inference test sessions with reconditioning sessions, extinction effects in response to the isolated auditory cues (*X_n)_* were minimised. At the end of each inference test day, mice were re-exposed to the observational learning stage (Figure 1c).

Reward-seeking behaviour was quantified by estimating the percentage of time spent in the area around the dispenser either during the 8 s visual cues (*Y_n,_* conditioning trials), or in the 10 s period after the auditory cues when no cue was presented (*X_n,_* inference test trials) (Figure 1d-e). The area around the dispenser was defined for each recording day using a 12 cm x 15 cm box, positioned such that the dispenser was in the centre of the 15 cm boundary, and the remainder of the box positioned within the open-field.

### Recording procedures

Following implantation surgery, mice recovered for at least two weeks before familiarisation to the recording procedure, with any calcium imaging recordings performed at least four weeks after viral injection. Mice were handled daily and exposed to the sleep box for >0.5 h per day for at least two days. During this period, mice were habituated to the miniature microscope mounting procedure and tetrodes were slowly lowered to the target locations. On the morning of each recording day, the electrophysiological profile of the local field potentials (LFPs) was assessed during sleep and used to adjust the position of each tetrode, to ensure the correct depth and maximise the cell yield. Tetrodes were left in position throughout the recording day. At the end of the recording day, tetrodes were raised (∼ 150 μm) to protect cell layers from potential mechanical damage overnight. We lowered each individual tetrode the next morning, making it unlikely that the recorded units were the same neurons across days.

Recordings were acquired on all days of the inference test (Figure 1a, 1c), using either calcium imaging data combined with electrophysiological recordings (n=4), or just electrophysiological recordings (n=5). To track the location of the animal, three differently coloured LEDs were attached to the microdrive casing on the mouse’s head and connected to the Intan board and images captured at a sampling rate of 39 Hz using an overhead camera together with the PosiTrack software (https://github.com/kevin-allen/positrack/wiki). At the end of each recording day, the mice were placed in a sleep box containing sawdust bedding and nesting material (12 x 12 x 28 cm, length x width x height) to record neural activity during sleep. Each open-field or sleep box recording session lasted ∼15-30 mins. Experiments were performed under dim light conditions (∼20 lux), with low-level continuous background white noise played through the speakers (except when auditory cues *X_n_* were delivered).

#### Electrophysiology recordings

Electrophysiology recordings were acquired from the electrodes and were amplified, multiplexed, and digitised using a single integrated circuit (headstage) located on the head of the animal (RHD2164, Intan Technologies, USA; http://intantech.com/products_RHD2000.html). The amplified and filtered (pass band 0.09 Hz to 7.60 kHz) electrophysiological signals were digitized at 20 kHz (RHD2000 Evaluation Board) and saved to disk with the synchronization signals from the positional tracking, the presentation of each type of sensory cue, the delivery and removal of the outcome drops, the lick events, and the (optional) calcium imaging acquisition.

#### Calcium imaging acquisition

Calcium imaging data were recorded using a miniature epifluorescence microscope (Inscopix nVista 3.0 Imaging System for In Vivo Research), attached above the prism lens implanted in V1 where cells were labelled with GCaMP6f, together with nVista (Inscopix) data acquisition box and acquisition software, using the following settings: 900 μm x 650 μm; Gain: 3, LED Power: 30-40%; sampling rate: 20Hz.

### Data pre-processing

#### Electrophysiology preprocessing

Spike sorting and unit isolation were performed via the automated clustering software, Kilosort (https://github.com/cortex-lab/KiloSort)^75^. To apply KiloSort to data acquired using tetrodes, the algorithm restricted templates to channels within a given tetrode bundle while masking all other recording channels. The resulting clusters were manually curated to check all clusters and remove spurious units using metrics derived from the cross-channel spike waveforms, auto-correlation histograms and cross-correlation histograms within the SpikeForest framework (https://github.com/flatironinstitute/spikeforest)^76^. All sessions recorded on a given day were concatenated and cluster cut together to monitor cells throughout the day. Each unit used for analyses showed consistent spike waveforms and stable firing rates throughout the entire recording day.

#### Calcium imaging preprocessing

Data acquired using calcium imaging were processed using Inscopix Data Processing software (1.6.0). Sessions were either combined into time series (if the miniscope was not disconnected between sessions) or preprocessed individually. Videos were cropped, spatially down-sampled by a factor of 2, bandpass filtered (0.005 to 0.5 pixel^-1^), and motion-corrected. Pixel values were normalised using a ΔF/F algorithm which took the mean image as a baseline. To identify individual cells, principal component analysis (PCA) and independent component analysis (ICA) were applied to the preprocessed data. This identified up to 150 independent components (ICs) per time series/session. We first applied automated sorting (cell size: 7-70 pixels; signal-to-noise ratio: 3; event rate: 0.0001Hz). We then manually checked the ICs to detect and remove spurious cells. We used longitudinal registration (minimum correlation: 0.5) to combine each session from a given recording day and only accepted cells that showed activity across all segments included in the registration.

For each accepted cell, calcium fluorescence traces were extracted from the corresponding spatial footprint generated by the PCA/ICA algorithm. Fluorescence traces were expressed as ΔF/F to give calcium transients and were exported for subsequent analyses. Moreover, calcium event trains were identified using the Inscopix event detection algorithm, which detects statistically significant increases in fluorescence relative to the local baseline while accounting for noise in the signal. Detected events were represented as binary timestamps. Only events that exceeded the software’s default significance threshold were included in downstream analyses.

### Local field potential signals

LFP signals were processed by first applying an anti-aliasing filter (8^th^-order Chebyshev type I filter) to the wide band signals sampled at 20 kHz. These signals were then down-sampled to 1,250 Hz using the decimate function from the signal submodule of SciPy (version 1.11.2).

### Sharp-wave ripple detection

For the LFP of each dCA1 tetrode positioned in the pyramidal cell layer, we subtracted the mean across all channels (common average reference), and applied dual stage-filtering using a band-pass filtered for the ripple band (80-250 Hz; 4^th^ order Butterworth filter) and a high-frequency bandpass filter (200-500 Hz, 4^th^-order Butterworth Filter). Instantaneous signal characteristics, including envelopes (instantaneous amplitudes), were derived using the Hilbert transform. Ripple events were identified by detecting envelope peaks within the ripple band that exceeded a threshold of five times the median value. In instances of multiple peaks within a 20-ms window, only the peak with the highest amplitude was considered. For each event we then identified its onset and offset points as the points where the ripple envelope fell below half of the detection threshold. Finally, we validated candidate ripple events using four criteria: *(i)* Ripple band power in the event channel, calculated as the squared mean amplitude, was at least twice the ripple band power in the common average reference (to eliminate common high frequency noise); (ii) The mean frequency of the detected event exceeded 80 Hz. (iii) Each event had at least four complete ripple cycles (to eliminate events that were too brief); (iv) Ripple band power was at least two times higher than the supra-ripple band defined as 200–500 Hz (to eliminate high-frequency noise, not spectrally compact at the ripple band, such as spike leakage artefacts). For events passing these criteria, the local maximum of each envelope was taken as the peak of the ripple, and these timestamps were recorded. On recording days with several tetrodes in the CA1 pyramidal layer, we used the tetrode with the largest mean ripple envelope amplitude to detect ripple events. We focused this study on ripples detected during sleep epochs, where all detected ripples were included in analyses unless otherwise stated.

## Quantification and statistical analyses

### Classifier analysis

Logistic regression classifiers were used to test whether behavioural choice (Figure 3a) and task cues (Figure 3b, Figure 6b) could be decoded from neural activity. The logistic regression classifier was implemented using the Python package *scikit-learn (1.6.1)*, with training and testing data split using stratified k-fold cross-validation, where the number of splits is set to the minimum number of cue presentations minus one. The classifier was trained on population vectors (across neurons) indicating the mean z-scored activity (V1: calcium transients; dCA1: spiking activity) across task cues. To decode subsequent behavioural choice, activity during auditory cues (*X_n,_* first 5 s) in the inference test was used to construct the population vectors. To decode task cues, activity during all six task cues was used to construct the population vectors (i.e. *X_1,_ Y_1,_ Z_1,_ X_2,_ Y_2,_ Z_2,_* full cue period). At least one presentation of each task cue was included in each fold and final decoding accuracy was calculated as the mean across all test folds. Data from any given recording day were only included if there were >3 correct responses for all relevant cues included in the test set.

### Representational similarity analysis

To assess the representational similarity of V1 ensemble neurons in response to the six task cues (*X_n,_ Y_n,_ Z_n)_*, we calculated a set of population vectors across V1 neurons. Each population vector corresponded to a single cue presentation (i.e. a single trial) and indicated the mean z-scored calcium transients across the duration of cue presentation (Figure 3c). For each recording day, we then constructed a representational similarity matrix (RSM) using the Pearson correlation coefficients between each pair of population vectors (“Data RSM”, Figure 3c-d). Next, we obtained the Pearson correlation coefficient between the “Data RSM” and a “Model RSM” (Figure 3e-h). The distributions of these correlation coefficients were then compared against a null distribution generated using permutation testing.

To test evidence for discrimination of set 1 and set 2 cues within a single modality (e.g. *X_1v_*s. *X_2)_*, we calculated a “Data RSM” for each cue type (*X_n,_ Y_n,_ Z_n)_* (e.g. Figure 3d for example showing *X_n)_*, before correlating the “Data RSM” with a “Model RSM” that included high correlation coefficients within-set and low correlation coefficients across-set (Figure 3e-f). A positive correlation between this “Data RSM” and “Model RSM” suggests that population activity in V1 distinguishes between set 1 and set 2 cues of the same modality (e.g. *X_1_* vs *X_2)_*.

To test evidence for predictive neural codes (e.g. *X_1_* → *Y_1;_ X_2_* → *Y_2)_*, we calculated a “Data RSM” by correlating population vectors that derive from different modality cues (*X_n_* vs. *Y_n)_*, and then correlated the “Data RSM” with a “Model RSM” that included high correlation coefficients within-set, and low correlation coefficients across-set (Figure 3g). A separate “Data RSM” was constructed for correct and incorrect trials, where for set 1 correct trials were defined as trials where two conditions were met: *(i)* the mouse visited the dispenser in the 10s following the cue; *(ii)* the mouse’s speed at the dispenser was less than 1 cm/s. All other set 1 trials where conditions *(i)* and *(ii)* were not met were classified as incorrect. The reverse was true for trials from set 2, such that incorrect trials were defined as those where conditions *(i)* and *(ii)* were met.

To control for the effect of behavioural state and spatial information, we used a GLM to further test evidence for predictive codes (e.g. *X_1_* → *Y_1;_ X_2_* → *Y_2)_* in our “Data RSM” (Supplementary Figure 2). The GLM included three explanatory variables: *(1)* “Behavioural state”: pairs of trials were highly correlated if the mouse made spatial trajectories towards the dispenser on both trials, while all other pairs of trials had zero correlation. This explanatory variable was included to control for representation overlap in spatial trajectories leading to the dispenser while also explaining variance attributed to heightened arousal associated with reward-seeking behaviour. *(2) “*Task specific behaviour”: pairs of trials where the presented cue derived from the same set were highly correlated, while pairs of trials where cues derived from different sets were negatively correlated. Additionally, pairs of trials where at least one trial was incorrect had zero correlation rather than the correlation defined from the relationship of the cues. *(3)* “Spatial”: the z-scored Pearson correlation coefficients were estimated between the spatial trajectories of the mouse on each pair of trials. The spatial trajectories were calculated by binning the environment into 5 cm x 5 cm bins and calculating the amount of time the mouse spent in each bin on each trial (Supplementary Figure 2a-b). For each recording day, the beta coefficients from the GLM were plotted (Supplementary Figure 2c). The distributions of these beta coefficients were then compared against a null distribution generated using permutation testing.

### Identifying ripple modulated cells

To identify ripple modulated cells (Figure 4b-d) we assessed the response of each cell/unit to ripple events and compared this response to a control distribution. A unique control distribution was generated for each cell/unit by applying a circular shift to the cell’s neural activity across a total of 1000 shuffles. By applying this circular shift we aimed to disrupt the relationship between ripples and spikes/calcium transients, while preserving the temporal relationships between spikes/calcium transients. To implement the circular shift, a randomly generated offset, *N*, was generated for each shuffle and applied to the spike train or calcium transients. *N* was generated with the Python package numpy’s *random.randint* function (between 2*sample rate, inclusive, and 60*sample rate, exclusive), before being applied as an offset to the spikes/calcium transients in a rolling manner using numpy’s *roll* function, with *N* elements at the end of matrix being reintroduced at the start of the spike train/calcium transients.

For each cell/unit in both the real and shuffled data, we calculated the mean over all ripples of the z-scored response (firing rate/calcium transients) during a 400 ms window centred on peak ripple power. Across this 400 ms window, we then calculated the mean (ephys) or maximum (ca) of the mean z-scored response for each cell/unit, to quantify the ripple response. We then compared the “true” ripple response against the control distribution for the same cell/unit. Cells/units were considered to be significantly modulated by ripples if their “true” response was greater than the 95^th^ percentile of the control distribution generated over the 1000 shuffles for the same cell/unit. Recording days were included in this analysis if the total number of ripple events detected in the sleep session was >100.

### Assessing shared information content across brain regions

A cross-validated GLM analysis was used to predict the activity of single cells/units recorded using electrophysiology during ripples using the population activity of another brain region, at different or the same time points relative to the peak ripple power (Figure 4e-i). For each target cell/unit, we constructed a vector of the total number of spikes within a 100 ms window centred on the peak ripple power of each ripple. A GLM was then trained to predict this target vector, using the ensemble activity within a given time window from all cells in the same region as the target cell/unit (excluding the target cell/unit itself) and from all cells in the “prediction” region of interest (Figure 4e-f). The time windows were 100 ms in size with centres 20 ms apart, spanning a 400 ms window centred on the peak ripple power. For each day, the cross-validated GLM analysis was only run if there was ≥1 target cell/unit to predict and >2 cells/units in the predictor region. For any given recording day, all cells/units were included where the total spike count for the cell/unit across ripples was ≥200 spikes.

The cross-validated GLM analysis was implemented by randomly splitting the data (across ripples) into 10 equally sized cross-validation folds. For each fold, we used 9 out of the 10 folds to train the GLM, while the remaining fold was used for testing. For testing, the *prediction score* was defined as the Pearson correlation between the predicted scores and the actual scores. The prediction scores were then averaged across all 10 folds.

To generate a null distribution, the original associations between spike times and neuron identities were randomly shuffled. This surrogate version preserved the timing of each spike (and thus the population rate within each ripple) and the number of spikes aoributed to each neuron (and thus the firing rate of each neuron). This shuffling controlled for the overall increase in multi-unit activity during ripples, ensuring that prediction scores were not artificially inflated by elevated spike counts in the ripple window. Moreover, this shuffle was applied only to spikes in the “prediction” region of interest (Figure 4f), thereby preserving the population activity of the target region in both the real model and the shuffle control, to test whether the addition of the “prediction” brain region improved the prediction score, over and above activity in the target region itself.

For each cell we then estimated the *prediction gain*, defined as the difference between the prediction score for the actual data and the mean of the shuffled control distribution (see Figure 4g-h, *upper,* for mean **±** SEM) and estimated a p-value using the proportion of shuffled scores that were greater than or equal to the true predictive score. Across cells, we used a binomial test to assess whether the proportion of cells with p<0.05 could be expected above chance, where alpha was set to 0.05, or 0.05/21 to provide Bonferroni correction across time bins (Figure 4g-h, lower).

### Defining “cue” cells

To identify cue modulated cells (Figure 5), we used a general linear model (GLM) to select neurons that represented the different cues in the task (*X_1,_ X_2,_ Y_1,_ Y_2,_ Z_1,_ Z_2)_* (Figure 5a). To avoid biasing the analysis toward highly active neurons, we first calculated the z-scored calcium transient or the z-scored firing rate for each neuron, for each recording day. For each neuron, the z-scored activity vector, recorded during the inference task, was regressed onto a GLM which indicated which cue was presented. For the drop trials, we took the presentation period as the time during which the mouse was near the dispenser and the drop was available. This provided a β weight for each neuron across all 6 cues (Figure 5a). These β weights indicate how much the activity of a given neuron changes in response to each sensory cue. We defined “cue” cells as neurons with a positive β weight more than 2 standard deviations from the mean β weight across all neurons for that cue.

### Quantifying the temporal relationship between cue cells

To characterise the temporal dynamics of cue cells across the dCA1→RSCg→V1 circuit, we analysed the inter-spike interval between pairs of cue cells, where one cue cell in the pair derived from dCA1 (Figure 5f). For each cross-regional pair of cells, we calculated the mean difference in spike times across ripples, and used a binomial test to test evidence for spikes in RSCg and V1 firing after those in dCA1 cell pairs. Cells that were significant for several cues were considered only once (i.e. each cell pair is unique).

### Assessing for sequential activity using Temporal Delayed Linear Modelling

We used Temporal Delayed Linear Modelling (TDLM)^53^, a domain general method, to test evidence for temporal structure in the reactivation of task cues during periods of sleep in the calcium transients recorded from V1 (Figure 6). Specifically, we tested evidence for neural sequences that respected the learned and inferred transitions in the inference task (*X_n_* → *Y_n;_ Y_n_* → *Z_n;_ X_n_* → *Z_n)_*.

First, we used spike trains (dCA1) or calcium transients (V1) recorded during the task to train a cross-validated logistic regression classifier (as described above) to predict the identity of the cue presented. For the V1 calcium data, we then applied the logistic regression classifier to data acquired in the sleep session to generate a timecourse of reactivation probabilities for each cue, generating a state space with dimension of time by cues. TDLM was then applied using a two-step process to identify sequences of memory reactivation during the sleep session. In the first step, we assessed the temporal relationship between each possible pair of cues *C_i_* and *C_j_*, by finding the linear relationship between the *C_i_* time series and the Δ*t*-shifted *C_j_* time series. Together, these *n*^2^ relations comprise an empirical transition matrix for each lag of Δ*t*, describing how likely reactivation of each cue is to be succeeded at a lag of Δ*t* by reactivation of each other cue. In the second step, we linearly related this empirical transition matrix for each lag of Δ*t* to a hypothesised transition matrix that reflects the task structure (Figure 6f). This produced a single number (which we refer to as “sequenceness”) that quantified the extent to which the neural data reflected the hypothesised transitions at any given lag of Δ*t*. Finally, we repeated this entire process for all Δ*t* of interest, yielding a measure of “sequenceness” at each possible lag.

To control for autocorrelations and neuronal overlap in the representation of different cues within a given set, we further estimated the “sequenceness” metric as the difference between evidence for forward (fwd) and backward (bwd) sequences, where the bwd “sequenceness” metric was generated by transposing the hypothesised transition matrix and re-estimating the sequenceness metric.

To test statistical significance we compared the final “sequenceness” metric against a null distribution generated by shuffling the hypothesised transitions matrix 10,000 times and in each shuffle estimating the final “sequenceness” metric. To define the two-tailed significance we took the 2.5^th^ and 97.5^th^ percentiles of the null distribution. To control for multiple comparisons across time lags, the statistical significance threshold was set to the minimum/maximum of the 2.5^th^/97^th^ percentile across all possible time lags. Over the course of the sleep session, using the optimal time lag from the TDLM analysis, we obtained a z-scored “sequenceness” score reflecting evidence for “sequenceness” at each time point. For each recording day, we then estimated the difference between the mean z-scored “sequenceness” within ripple windows (400 ms centred on the peak ripple power) and a control distribution. The control distribution was generated by estimating “sequenceness” at random time points, where the random time points were selected by calculating the time interval between each ripple and randomly shuffling these time intervals to extract “sequenceness” metrics at permuted ripple timepoints, across 10,000 permutations (Figure 6n).

### Quantifying co-activity between triplets and doublets of cue cells during ripples

To determine whether triplets or doublets of cue cells in V1 were coactive during ripples (Figure 7a-f), for each recording day we estimated the probability of joint calcium event trains during ripples recorded during the sleep session. Recording days were included if the total number of ripple events detected in the sleep session was >30. For each ripple, the peri-ripple time window was defined as **`±`**1000 ms from peak ripple power, to allow sufficient time for triplet sequences of calcium event trains occurring at the optimal time lag estimated using TDLM (Figure 6g-h). Within these ripple windows we estimated the probability of joint calcium event trains for cue cells representing *X_n,_ Y_n_* and *Z_n._* To this end, we first listed all possible triplets, before computing the coactivation probability for each triplet as follows:

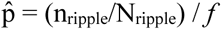

where n_ripple_ is the number of ripples during the sleep session where there were detected calcium events for all cue cells in the triplet; N_ripple_ is the total number of ripples during the sleep session for the recording day; *f* is the average of the mean firing rate of all cue cells in the triplet, across the sleep session. To assess evidence for representation of the within-set inferred relationships (*X_n,_ Z_n)_* in V1 during dCA1 ripples, we estimated the coactivation probability p^ of joint calcium event trains for cue cells representing the doublet *X_n_* and *Z_n,_* but only in absence of calcium event trains from cue cells representing *Y_n._* For both triplets and doublets, we then tested whether the mean coactivation probability p^ across recordings days was significantly different from a null distribution. The null distribution was constructed by permuting the ripple windows within the sleep session 1000 times. For each permutation, within the sleep session we randomly shuffled the time of peak ripple power for each ripple event, before taking the mean coactivation probability p^_co*nt*rol_ across recording days, as follows:

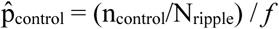

where n_co*nt*rol_ is the number of permuted ripples during the sleep session where there were detected calcium events for all cue cells in the triplet/doublet of interest.

For visualisation, of the coactivation probability for both triplets and doublets, for each triplet/doublet we estimated the difference, p^_diff_, between the coactivation probability p^ during ripples, and the mean of the coactivation probability p^_co*nt*rol_ across all permutations, as follows:

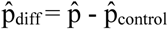

before using 10,000 bootstrapped resamples to illustrate the effect size (Figure 7b-e).

### Quantifying the directional relationship between pairs of cue cells

The temporal relationship between z-scored activity in pairs of cue cells across the sleep session were quantified using cross-correlograms (Figure 7g-j), applied either to calcium transients or electrophysiology data. In each case, the activity of the second cell in the pair was shifted while holding the activity of the first cell in the pair fixed, as reference, such that: negative time lags indicate backward (bwd) sequences; positive time lags indicate forward (fwd) sequences. The time windows for the cross-correlograms approximated 2x the optimal time lag estimated using TDLM, to allow for triplet sequences, corresponding to -5 ms to +5 ms for spike data and -400 ms to +400 ms for calcium data. Bootstrapping was used to illustrate the difference in means between fwd and bwd sequences. Permutation testing was used to test statistical evidence for fwd versus bwd sequences.

### Statistical plotting

Bootstrap-coupled estimation (DABEST) plots^37^ were employed to estimate effect sizes (Figures 1, 4–5, 7), computed from 10,000 bias-corrected bootstrapped resamples^38^. In each case we indicate the mean using a black dot, 95% confidence interval using black ticks, sampling-error distribution using a filled-curve.

### Permutation testing

We used permutation testing to test statistical significance. For each test, we generated a null distribution by shuffling the identity labels on the data before estimating a permuted test statistic of interest, across 10,000 iterations (unless otherwise stated). The observed test statistic was then assessed against this null distribution, with the p-value estimated as the proportion of permuted test statistics that are more extreme than the observed test statistic.

## Acknowledgments

We would like to thank B. Micklem and J. Westcott for technical support, Rafal Bogacz for inspiring discussion on the data, Leonie Glitz and Sumedha Nalluru for comments on a previous version of the manuscript and the Barron lab and Dupret lab for discussions at all stages of the project. This research was funded by the UKRI (MR/W008939/1 to H.C.B.), MRC (Program MC_UU_00003/4 and MR/W004860/1 to D.D.). C.M.S. was supported by the British Neuroscience Association Scholars Programme. The Medical Research Council Centre of Research Excellence in Restorative Neural Dynamics is funded by the MRC (UKRI/MR/B000936/1). The funders had no role in study design, data collection and analysis, decision to publish or preparation of the manuscript.

## Author contributions

All authors contributed to the preparation of the manuscript. P.V.P., D.D. and H.C.B. developed the methodology for multi-site electrophysiology and for calcium imaging; H.C.B. developed the methodology for combining electrophysiology and calcium imaging to allow for simultaneous acquisition; C.M.S., O.J.M. and H.C.B. acquired the in-vivo electrophysiology and calcium imaging data; P.V.P. and D.D. developed the trans-synaptic virus; C.M.S., M.C.C.G., O.J.M. and H.C.B. performed viral tract-tracking, including immunohistochemistry; C.M.S., M.C.C.G., O.J.M., R.M.H., H.C.B. acquired confocal images; C.M.S. analysed the data, with help and/or advice from V.LDS., M.C.C.G., R.M.H., D.D., H.C.B.; C.M.S., M.C.C.G., R.M.H. and H.C.B. wrote the paper; D.D. and H.C.B. acquired funding.

## Data availability

All data generated and analysed during this study are included in the manuscript and supporting files. Upon publication the data will be made available via Cambium, the sharing platform of the MRC CoRE in Restorative Neural Dynamics https://data.mrc.ox.ac.uk/data-set/ephys-ca-generative-replay.

## Code availability

Upon publication the code used in this study will be made available via Cambium, the sharing platform of the MRC CoRE in Restorative Neural Dynamics https://data.mrc.ox.ac.uk/data-set/ephys-ca-generative-replay-code.

