## Supplementary figures and images for "Generative replay across hippocampal-neocortical circuits"

### Supplementary Figure 1

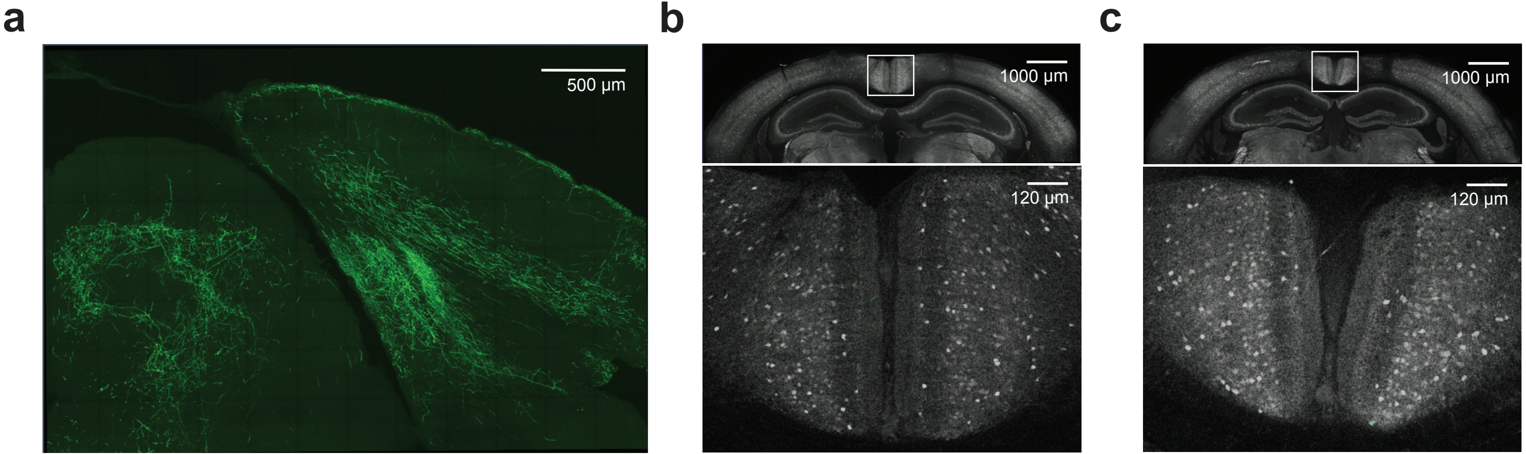

### Supplementary Figure 2

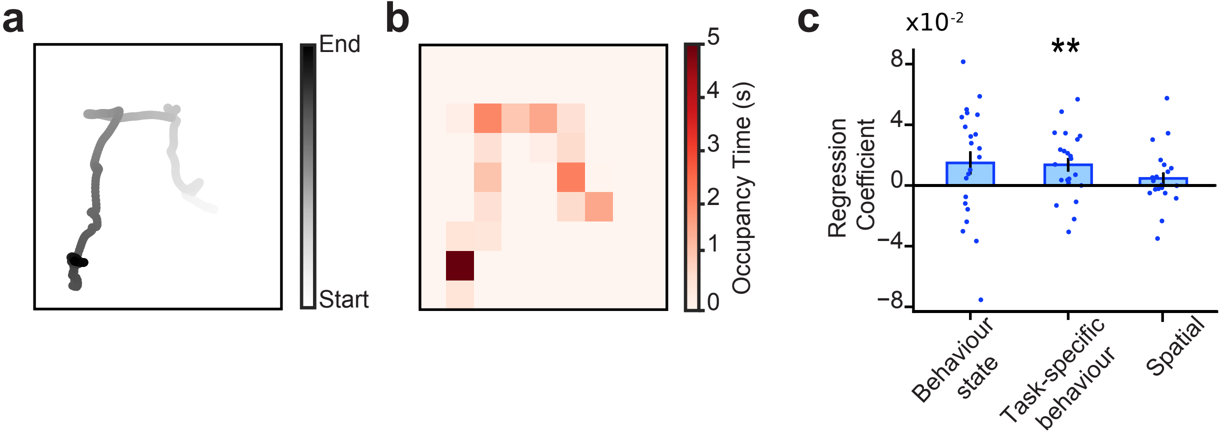
